# Emergence of consistent intra-individual locomotor patterns during zebrafish development

**DOI:** 10.1101/634451

**Authors:** Jennifer A. Fitzgerald, Krishna T. Kirla, Carl P. Zinner, Colette M. vom Berg

## Abstract

The analysis of larval zebrafish locomotor behavior has emerged as a powerful indicator of perturbations in the nervous system and is used in many fields of research, such as neuroscience, toxicology or drug discovery. The behavior of larval zebrafish, however, is highly variable, resulting in the use of high numbers of animals and the inability to detect small effects. In this study, we analyzed whether individual locomotor behavior is stable over development and whether behavioral parameters correlate with physiological and morphological features of the larvae, with the aim to better understand variability and predictability of larval locomotor behavior. We found that locomotor activity of individuals is consistent within the same day and becomes predictable during development especially during dark phases, when larvae are performing exploratory light-searching behavior and display increased activity. Stimulus induced startle responses were less predictable for an individual, and response strength did not correlate with inherent locomotor activity. Moreover, locomotor activity was not associated with physiological and morphological features of the larva (resting heart rate, body length, size of the swim bladder). These findings highlight the areas of intra-individual consistency, which could be used to improve the sensitivity of assays using zebrafish locomotor activity as an endpoint.

## Introduction

The ontogeny of zebrafish locomotor behavior and the underlying maturation of the locomotor network has been subject of extensive studies, fueled by the vast variety of genetic, molecular, physiological and behavioral tools developed for this prominent vertebrate model organism. The first embryonic movements start around 17 hours post fertilization, but it is not until 2-3 days post fertilization (dpf) that the larvae swim spontaneously^1,2^. This swim pattern is initially infrequent and in bursts, that slowly transitions into beat-and-glide swimming mode after swim bladder inflation and before feeding at 5 dpf ^3,4^. This sequence of events and the underlying cellular mechanisms are described in the literature^5–7^, and as a result, the analysis of zebrafish locomotor activity has become a popular read-out to assess the impact of external challenges to the nervous system in many fields of research. The amenability to high-throughput, non-invasive analysis, which allows cost-, material- and time-effective testing as compared with other vertebrate model organisms, additionally contributes to the popularity of zebrafish behavioral assays. Moreover, the availability of commercial plug and play systems (e.g. from Noldus, Viewpoint or Loligosystems) has facilitated behavior data acquisition and analysis, making this an endpoint that can now be readily used. These locomotor tests, however, suffer from high inter-individual variability and small but important effects can neither robustly nor repeatedly be detected^8–14^.

Behavioral inter-individual variability is common within populations of organisms, and the concept of individuality and personality has been reported for humans^15,16^, birds^17,18^, fish^19^, and other species^20,21^. Behavioral variability can arise from genetic, developmental, pharmacological, environmental and social processes^22,23^ and plays an essential role in the response and adaptation of a population to environmental changes^24–26^. Despite this importance, the variation among individuals is often ignored when behavior is quantified as averages with associated dispersions and individuals within a group are generally considered as simple replicates^27–30^. However, environmental changes can affect variation without changing the mean and potential biological significance is obscured when such variation is ignored. Therefore, it is important to address and understand these differences between individual behaviors as it could facilitate the understanding of an individual’s response during environmental adaption^25,26^.

However, inter-individual differences within a population are not the only issue that results in large variation within, especially behavioral, experiments. High variability within an individual’s own response can also contribute to variation, with these intra-individual differences mainly attributed to ontogenetic and environmental effects^31–33^. While intra-individual consistency in behavior has been widely addressed in primates and rodents, aquatic models are less characterized in this regard despite their increasing use in behavioral trials^19^. For the testing of acute effects on the nervous system, whether of toxicants, drugs, stressors or other perturbations, the existence of intra-individual consistency of locomotor behaviors in early larval zebrafish stages would allow baseline measures of locomotor activities of all individuals prior to exposure to which effects can then be normalized to. This would allow a better estimation of effects, especially if these are small, thereby increasing the sensitivity of such tests.

Therefore, the goal of this study was to test whether consistency of locomotor activity of an individual zebrafish emerges during larval development and under which conditions this may occur. Given that light conditions shape locomotor patterns differently^34–37^, we hypothesized that intra-individual consistency might vary under different light conditions. In addition, we tested whether consistency can be observed from stimulus-triggered activity responses and whether individual differences can be attributed to physiological or morphological features of the larva.

## Results

### Locomotor behavior is most predictable in darkness

To study the consistency of locomotor behavior of an individual larva over time, a total number of 132 mixed wildtype (WM) larvae were subjected to different behavior tests at two time points (9am and 2pm) over three consecutive days (5, 6 and 7 days post fertilization, dpf; Fig.1a). As the locomotor behavior of zebrafish larvae changes under different light conditions^34–37^, we analyzed spontaneous swimming after a 20 min period of acclimatization (referred to as “spontaneous”), swimming under darkness (2 × 10 min, referred to as “dark intervals”) and swimming in light after periods of darkness (2 × 10 min, referred to as “light intervals”) (Fig. 1a). Fig. 1a demonstrates short peaks of increased activity at the light switches as well as heightened locomotor activity during dark intervals. In addition, we investigated whether inherent locomotor activity of individual larvae relates to their activity during startle responses, triggered firstly through 4 one second dark flashes (Fig. 1b) and secondly by using a tapping stimulus device integrated in the behavior system^38^ (Fig. 1c). We chose an inter-stimulus interval of 90 seconds to measure the startle response from an individual repeatedly without inducing habituation. Short peaks of increased activity occurring immediately after stimulus application indicate that startle responses were triggered with these two protocols. The activity and radial index were measured to characterize the swimming behavior during the tests. The activity index is the percentage of time the individual moves within one-second intervals. The radial index indicates where the larva moves within the well and is calculated based on the distance of each larva in respect to the wall. Smaller indexes represent closeness to the wall and larger indexes represent more central locations. Although these parameters have been shown to be independent of each other^39^, for our data we can confirm this only for certain experimental protocols depending on when the experiment took place (Supp. Tab. 1).

**Figure 1.**
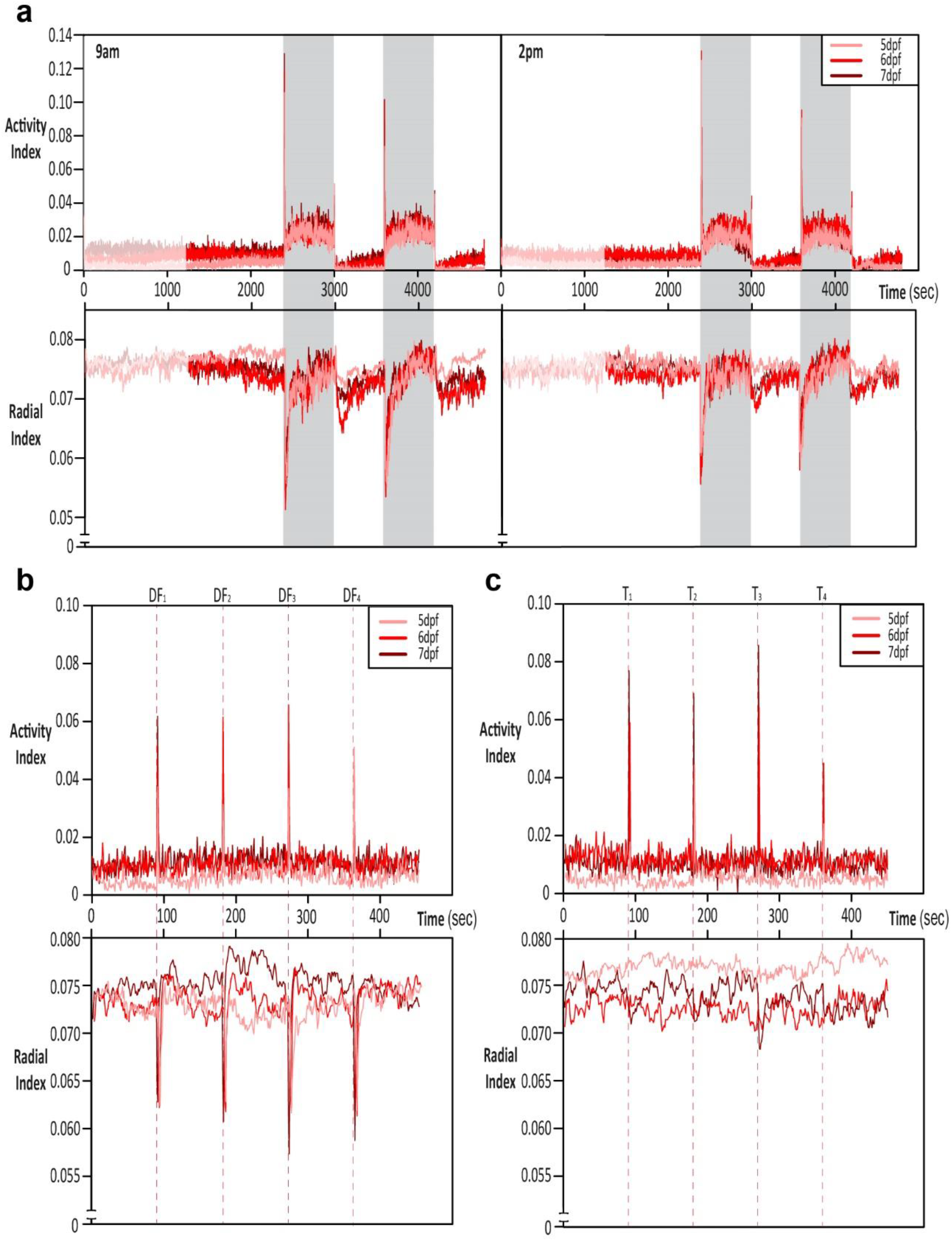
Time series plots from behavior experiments. a) Average activity index (top) and radial index (bottom) of 132 zebrafish larvae over the study period under light or dark period at either 9am or 2pm for 5, 6 or 7 dpf. Initial 20 minutes acclimation period (faded period), followed by 20 minutes of “spontaneous” swimming, then swimming under darkness (2 × 10 min, referred to as “dark intervals”, shaded period) and swimming in light after periods of darkness (2 × 10 min, referred to as “light intervals”). For the startle triggers of b) dark flashes (DF) and c) tapping (T) average activity for 9am measurements are shown. The dashed lines indicate occurrence of the stimuli.

We found that intra-individual variability was consistently low during dark intervals, while it was gradually decreasing over development during spontaneous swimming and light intervals (Fig. 2a). Differences in activity distribution could explain these changes, as the activity of fish is lower under spontaneous and light intervals compared to dark intervals. Most fish either do not move or display low activity during the spontaneous and light intervals while during dark intervals most fish display activity but the amount they move varies and is larvae dependent (Fig. 2b). This is further supported, as the lower coefficient of variation (CV) generally resulted from a higher mean activity rather than a smaller standard deviation (Supp. Fig. 1a - c), indicating that the more the larvae move, the less variable their movements are.

**Figure 2.**
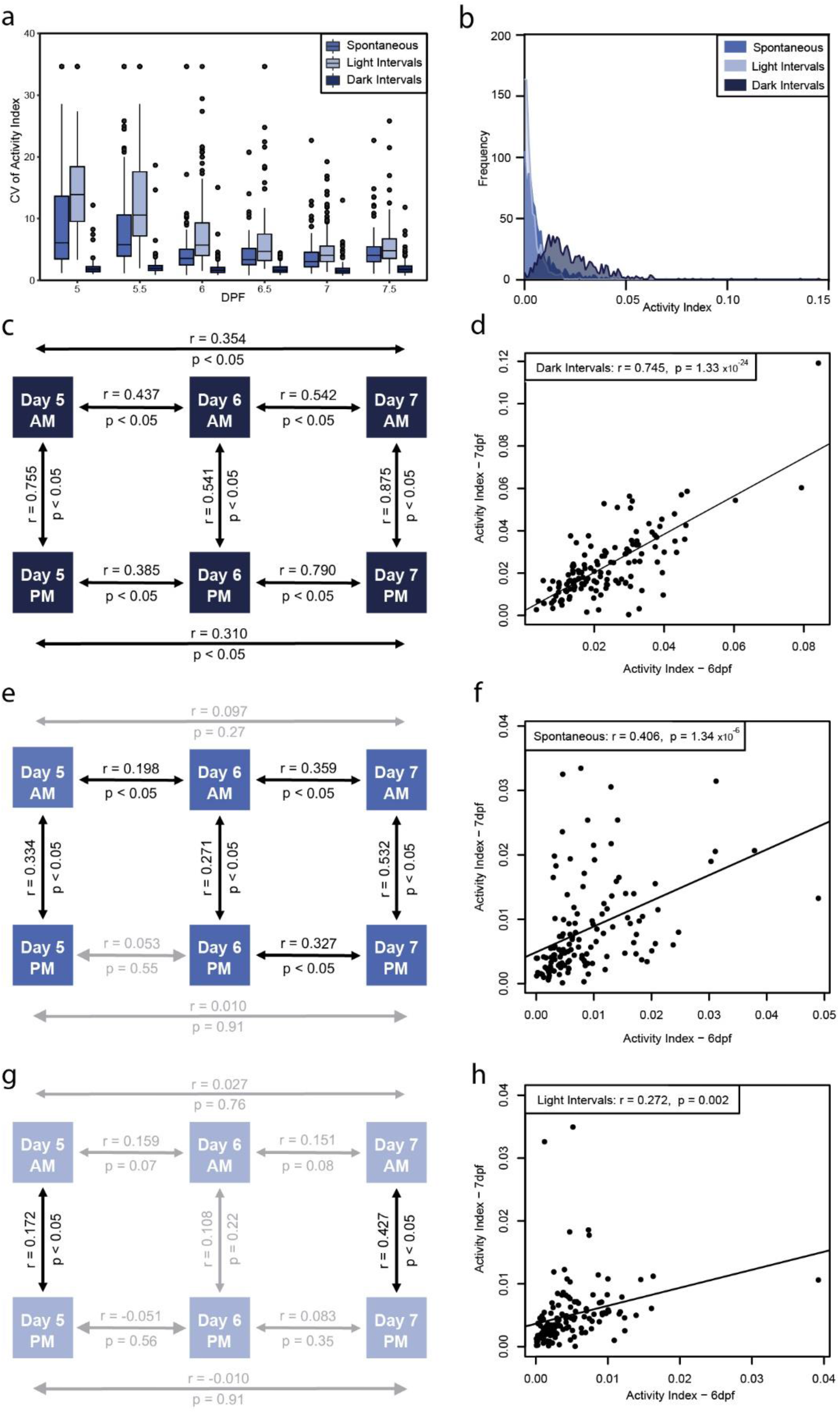
Behavioral intra-individual variability in a population of 132 larvae for the activity index. a) Boxplot of the coefficient of variance of activity index for each individual larva (n=132) over the different days and daytimes (days post fertilization; dpf) under the different conditions studied. b) Frequency distribution of the activity index in spontaneous, light and dark intervals depicting the differential activity profiles under these conditions. Schematics represent correlations of activity between different days and times of day for each of the conditions of study, c) dark intervals, e) spontaneous and g) light intervals. Correlation plots between activity of larvae on day 6 vs 7 for d) dark intervals, f) spontaneous and e) light intervals. Statistics on the plots represent the Pearson’s correlation coefficient and respective p value, with a linear regression line fitted for visual aid on the scatter plots.

Considering that the intra-individual variability was lowest during dark intervals, we analyzed whether an individual’s activity is consistently high or consistently low under dark conditions during development by looking at the correlation of activity between the different days (5, 6, 7 dpf) and time of day (9am and 2pm). There was a strong correlation between measurements taken at 9am compared to 2pm for all days measured (5 dpf: r = 0.755, p < 0.05; 6 dpf: r = 0.541, p < 0.05; 7 dpf: r = 0.875, p < 0.05; Fig. 2c), indicating that there was no effect of time of day on the larvae’s behavioral response to dark intervals. When looking at the different days, correlations were observed for all days (Fig. 2c), but the strongest correlations occurred between day 6 and 7 (r = 0.745, p < 0.05; Fig. 2d), suggesting that an inherent locomotor activity emerges at day 6. Interestingly, activity during spontaneous swimming was less predictable (Fig. 2e), although still showing a correlation, albeit weaker, between day 6 and 7 (r= 0.406, P < 0.05; Fig. 2f). In addition, for light intervals, the interactions were even more unpredictable (Fig. 2g): Most comparisons resulted in no correlation, and the correlation between day 6 and 7, although significant, was very weak (r = 0.272, p < 0.05; Fig. 2h), potentially as a result of the higher variance and lack of activity the fish displayed (Fig. 2a and b).

To see if the larvae can adapt to the experimental procedure, we additionally tested whether the correlations improved by starting the test on day 4. Due to the larvae’s lack of activity at 4 dpf, no significant trends could be determined when comparing day 4 to the other days, for any of the periods measured (Supp. Fig. 2). For light intervals, a slight increase in the strength of the correlations was observed, especially for 2pm measurements (Supp. Fig. 2), potentially due to a reduction in variation in the data that is observed from a high 5 dpf CV in the 3 day experiment.

When considering the location of the larvae in the well, less intra-individual variability occurs between the different experimental conditions tested, compared to the activity index (Fig. 3a). This indicates that larvae move very consistently, either swimming close to the wall or in the middle of the well, independent of the light conditions. Interestingly, between the different days and time points there is a larger proportion of significant interactions (Fig. 3e, f and g), albeit the actual strength of correlation tends to be weaker for the radial index compared to the activity index (e.g. for dark intervals- radial index: r = 0.694, p < 0.05; Fig. 3b, activity index: r=0.745, p < 0.05; Fig. 2d). This demonstrates that the radial index was less able to predict the movement between the two days compared to the activity index.

**Figure 3.**
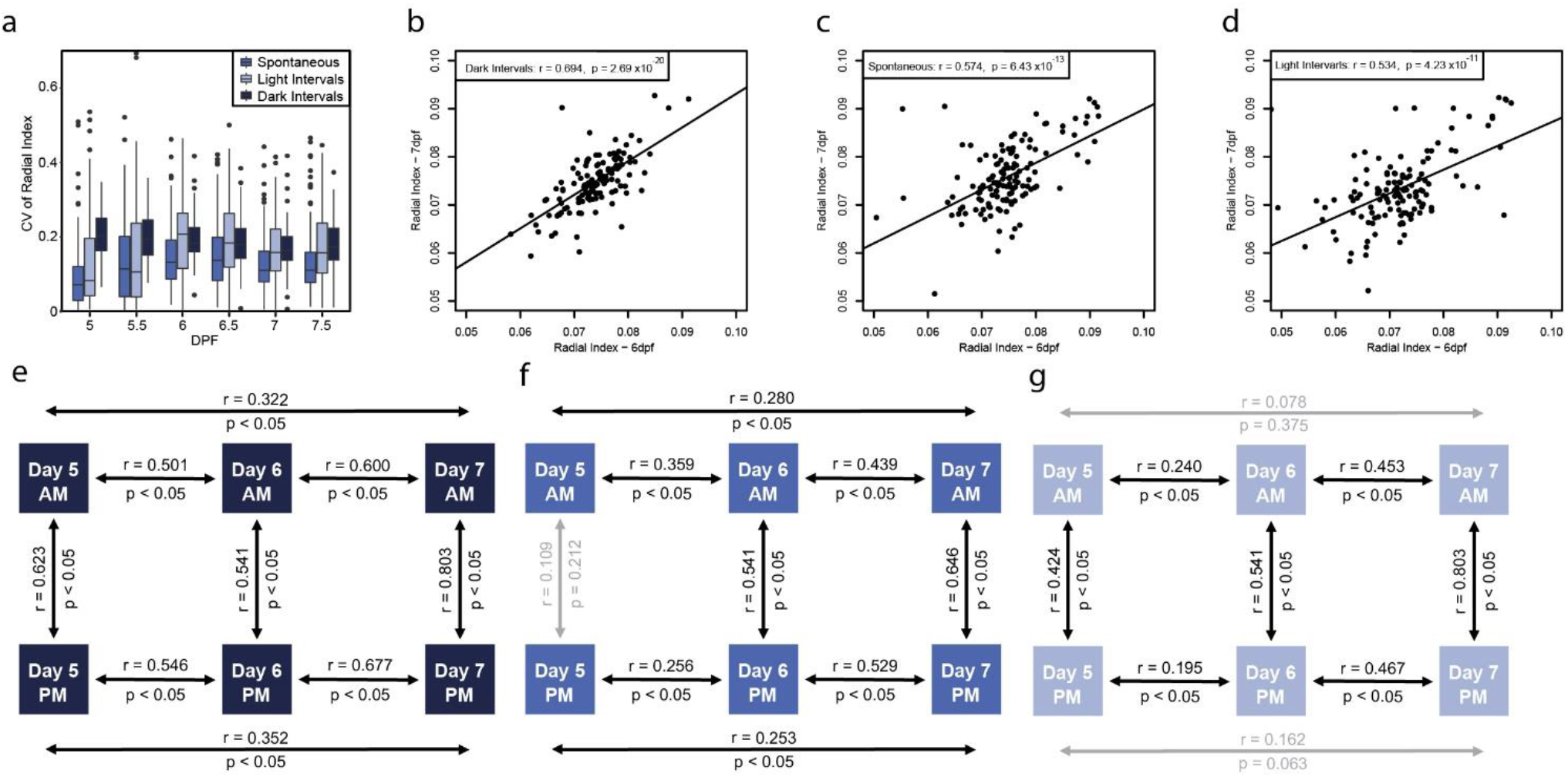
Behavioral intra-individual variability in a population of 132 larvae for the radial index. a) Boxplot of the coefficient of variance of radial index for each individual larva (n=132) over the different days and daytimes (days post fertilization; dpf) under the different conditions studied. Correlation plots between the radial index of larvae on day 6 vs 7 for b) dark intervals, c) spontaneous and d) light intervals. Schematics represent correlations of the radial index between different days and times of day for each of the conditions of study, e) dark intervals, f) spontaneous and g) light intervals. Statistics on the plots represent the Pearson’s correlation coefficient and respective p value, with a linear regression line fitted for visual aid on the scatter plots.

In summary, we show that although the activity patterns slightly differ between mornings and afternoons, individual zebrafish larvae move consistently, thus individuals with a high activity level in the morning also have a high activity level in the afternoon. Moreover, in darkness, when zebrafish larvae show hyperactivity compared to light conditions (Fig. 1a), intra-individual activity and radial index are most consistent and become highly predictable from day 6.

### Startle responses are not consistent for an individual

We tested whether inherent locomotor activity of individual larvae related to activity during a startle response, triggered from two different protocols (“dark flash” and “tapping”) (Fig. 1b and 1c), as well as induced when switching the lights on (“onset”) and off (“offset”) (Fig 1a). The strength of the startle responses was measured by calculating the distance moved during one second after the stimulus was applied. The average responses to each trigger showed some differences, however, no clear trend is discernible (Fig. 4a). In addition, the individual responses seem to show neither a consistency nor a uniform decrease in strength over the different stimuli, for both tapping (Fig. 4b) and dark flashes (Supp. Fig. 3), suggesting that fish are not consistently habituating to the stimuli. Indeed, when calculating a habituation index (HI) over the 4 stimuli, habituation occasionally occurs, but is not consistent for an individual over development (Supp. Fig. 4).

**Figure 4.**
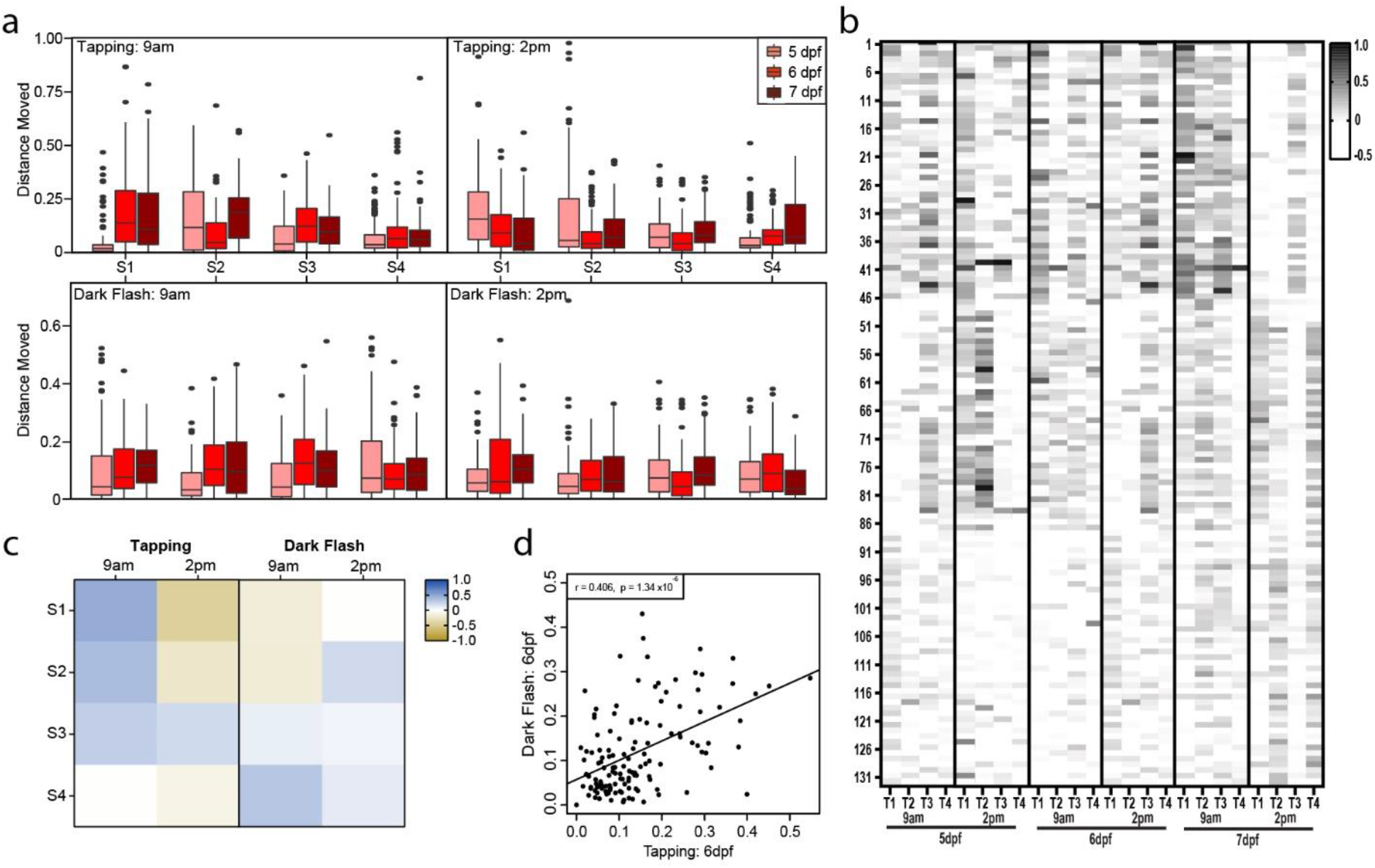
Individual responses to startle stimulus. a) Boxplots of the average distance moved of the 132 larvae at each of the stimulus (S1-S4) (tap or dark flash) at either 9am or 2pm. The different color boxes represent the three different days measured on 5, 6 and 7 days post fertilization (dpf). b) Heat map representing the change in distance moved with respect to the baseline of each individual larvae for all time points and days measured in response to the each of the tapping stimulus (T1-T4). White represents no response to the stimulus, with the grey scale darkening in a linear scale depending on the strength of the response. c) Heat map of the r values from the correlations of distance moved of individual larvae between 6 and 7 dpf, for each of the 4 startle stimuli at either 9am or 2pm for tapping and dark flashes. Blue represents a positive correlation, with yellow representing a negative correlation. d) Correlation plot between the individual fish response to the first dark flash and the first tap at 6 dpf. Each point on the graph represents an individual larva (n=132) and the correlation coefficient was calculated using Pearson’s correlation, with a linear regression line fitted for visual aid.

Looking closer at individual consistency, we found a significant, albeit only moderate correlation when comparing the response from the first tapping stimulus between 6 and 7 dpf at 9am (r = 0.423, p < 0.05; Fig. 4c). However, for responses in the afternoon this correlation becomes negative (r = −0.404, p < 0.05; Fig. 4c), potentially due to the larvae reducing their activity at 2pm compared to 9am which is supported by the comparison of the average distance moved between those periods (7 dpf 9am average distance moved: 0.138 ± 0.20; 7 dpf 2pm average distance moved: 0.041 ± 0.11; p < 0.05; Fig. 4a). The significance of this correlation weakens with each tapping stimulus and is ultimately lost (Fig. 4c). Interestingly, this trend was not seen at all for the dark flash stimulus, where no significant correlations between the response of the fish on day 6 to 7 for either 9am or 2pm, irrelevant of the stimulus number, was detected (Fig. 4c). The same inconsistency was found for responses triggered at onset and offset of light switches (Supp. Tab. 2).

When cross-comparing responses to tapping stimuli and dark flashes, we found a weak correlation for the first stimulus at day 6 (r = 0.406, p < 0.05; Fig. 4d). However, this was not consistently seen for all days or time points studied (Supp. Tab. 2), supporting further that the larvae’s response to a startle stimulus is not predictable or consistent at the individual or population level. Moreover, when comparing an individual’s inherent locomotor activity with startle response strength, no strong correlations were found for the different protocols used under all light conditions (Supp. Tab. 3), suggesting that the beat-and-glide swim mode does not relate to an individual’s startle response capabilities, irrespective of the stimulus modality.

### Locomotor activity is not associated with physiology and morphology

Resting heart rates in individual larvae can vary. We tested if this property links to their inherent locomotor activity. The resting heart rates of our WM larvae lies within a broad range of 118.60 – 225.46 beats per minute at 5 dpf with a mean of 185.55 BPM (± 21.94). The mean did not significantly change at 6 dpf (mean = 186.51 ± 16.17), but at 7 dpf there was a significant decrease compared to day 5 and 6 (mean = 176.63 ± 21.58, p < 0.05) (Fig. 5a). Despite this, the resting heart rate of an individual was significantly consistent over the three days measured (Fig. 5b, Supp. Tab. 4). However, although the resting heart rate and locomotor activity for an individual are consistent during development, they did not correlate with activity during dark intervals (5 dpf: r = 0.146, p = 0.10; 6 dpf: r = −0.015, p = 0.86; 7 dpf: r = 0.084, p = 0.34; Fig. 5c) or for the other light conditions tested (Supp. Fig. 5a and b).

**Figure 5.**
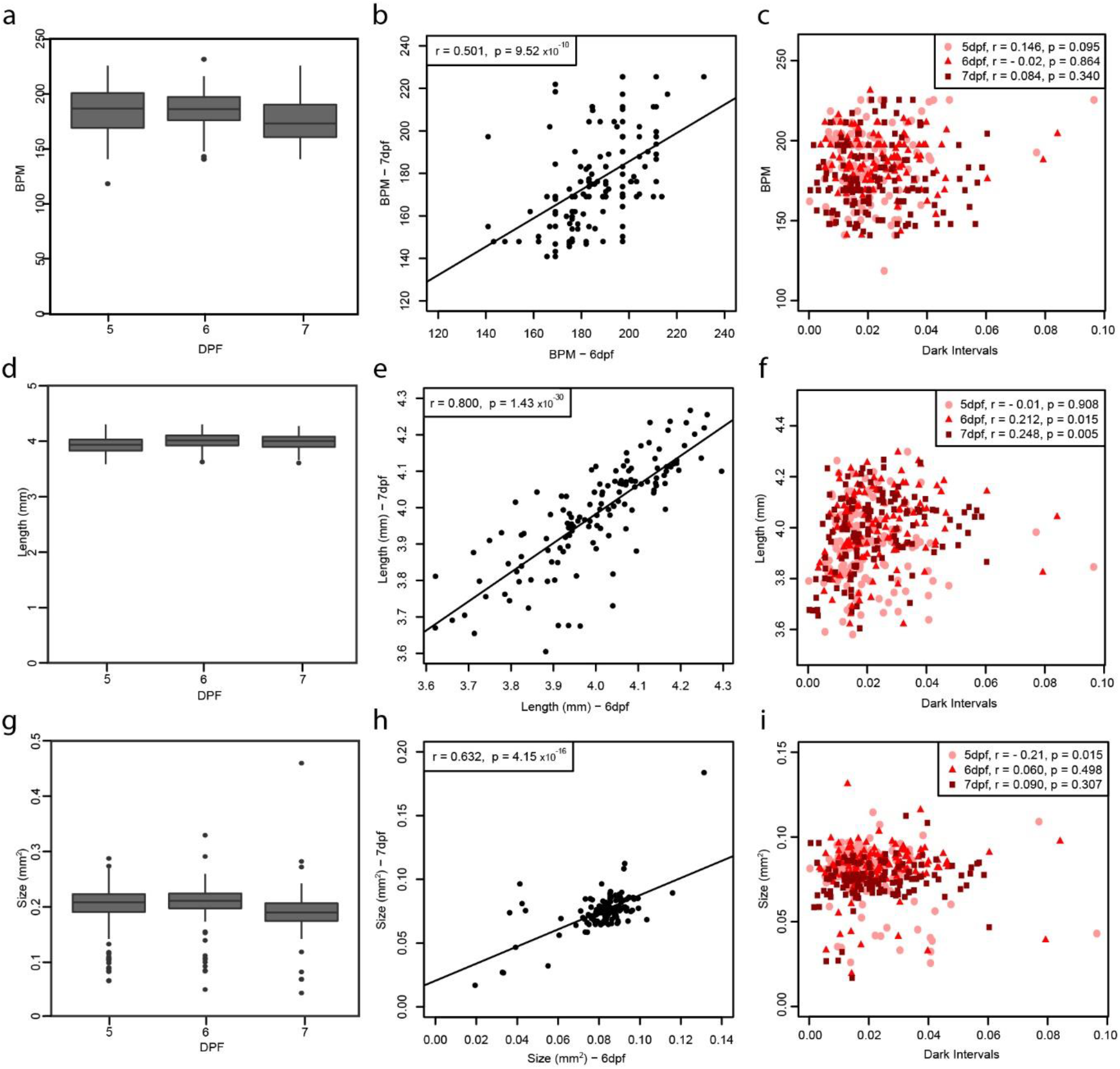
Physiology and morphometric parameter comparisons. Boxplots representing the average measure of a) heart rate (beats per minute; BPM), d) body length (mm) and g) size of swim bladder (mm^2^), over the three days of experiments (5, 6 and 7 days post fertilization, dpf) with significant difference represented by different letters on each graph. Correlation plots between 6 and 7 dpf for b) heart rate, e) body length and h) size of swim bladder, with each point representing a single larva. Comparison of the individuals’ c) heart rate, f) body length and i) size of swim bladder to their respective average activity during dark intervals, with each day plotted on each plot (5 dpf: circle, 6 dpf; triangle and 7 dpf; square). Statistics on the plots represent the Pearson’s correlation coefficient and respective p value, with a linear regression line fitted for visual aid on the scatter plots.

We further assessed morphological features of the larvae and tested whether they are consistent over time for each individual and whether they would underlie the differences in locomotor activity among the individuals. Body length was significantly smaller at day 5 compared to the other days, but there was no significant increase from day 6 to 7 (Fig. 5d), with size strongly correlated between the days tested, suggesting that the individuals grow at a constant pace (Fig. 5e, Supp. Tab. 4). Again during dark intervals, there was no strong correlation between activity of the larvae and their body length (Fig. 5f), with similar patterns observed for spontaneous movement and light intervals (Supp. Fig. 5c and d). Similarly, for the size of the swim bladder, fish that had a large swim bladder on day 5 consistently had larger swim bladders on day 6 and 7 (Fig. 5h, Supp. Table 4). However, on average, swim bladders appeared significantly smaller on day 7 compared to day 6 (Fig. 5g), potentially because of a change in shape, as the developing swim bladder undergoes a process to eventually form a double-chambered swim bladder in the adult stage^40,41^. There was no significant correlation between the activity of larvae under dark intervals and the size of the swim bladder, although for day 5 there was a slight trend towards fish with smaller swim bladders moving more (Fig. 5j), which was also seen for spontaneous movement and light intervals (Supp. Fig. 5e and f).

## Discussion

Behavioral diversity of a population can be observed throughout the animal kingdom in genetically diverse and even in isogenic populations^42,43^ and behavioral inter-individual variability as well as inter-strain variability is increasingly reported for laboratory strains of zebrafish kept for long periods of time^44–49^. Despite this, variation among individuals is often ignored when behavior is quantified as averages with associated distributions^24,27–30^. Therefore, in this study, we aimed to address the hypothesis that despite the high inter-individual variability in zebrafish locomotor activity, intra-individual consistency might emerge during larval development, possibly shaped by different light conditions and physical properties of the larvae.

Our data shows that locomotor activity begins to become predictable for an individual during development around 6 dpf, especially during darkness-induced explorative behavior. We also demonstrate that locomotor activity does not correlate with physiological and morphological features in larval stages, although these features are consistent within an individual. These conclusions are supported by previous research showing that swimming behavior is predictable between individuals when swimming freely in identical wells^39^. Roman *et al*. identified a histone H4 acetylation pathway that modulates individual behavior in a genetics-independent manner without affecting the global average behavior of the population. Therefore, while the average behavior might mostly depend on genetic background or environmental changes, behavioral inter-individual variability could result from histone H4 acetylation differences.

The most consistent period for all behavior parameters measured was between 6 and 7 dpf, with the most predictable activity under dark intervals, compared to spontaneous and light intervals. Spontaneous locomotion in zebrafish larvae has been shown to follow a non-random pattern even in the absence of sensory cues, facilitating the detection of resources or shelter^50^. Driven by the dwindling nutritional stock supplied from the yolk, the developing larva starts actively hunting for food from around 5 days post fertilization. This predation is strongly dependent on a functional visual system, as larvae in darkness or with impaired vision are unable to locate prey^51,52^. Accordingly, upon change of light conditions, larvae engage in different light-search behaviors to locate prey. These include phototaxis, where light is restricted to an area and movements towards the light source are guided by retinal input^53,54^, or dark photokinesis, where illumination is completely lost, and locomotion is strongly increased^34,35^ and largely driven by non-retinal deep-brain photoreceptors expressing the light-sensitive pigment melanopsin^36^. Recent findings, however, show that heightened locomotor activity of the larvae during darkness is not random and undirected, as implied by the definition of photokinesis, but rather structured and resembles an area-restricted local search in a first phase followed by a more outward-directed roaming search to efficiently detect light sources^37^. With increasing age, decreasing yolk and under dark conditions, search strategy behavior becomes more important for the larvae and likely causes individuals to unwind their hard-wired program. This possibly explains our finding that the intra-individual consistency is highest under dark conditions and with advanced larval age under unfed conditions.

We performed our tests during 5 to 7 days post fertilization, as we aimed to find conditions for consistency in a standard well plate and for ages that are frequently used for zebrafish behavior studies. In addition, we chose this period to be able to test under unfed conditions, to avoid introducing another variable through feeding behavior. Food can introduce confounding factors in a high-throughput testing set up for drugs, pollutants or other perturbations, so identifying specific predictable periods for behavior testing without food is preferable^55,56^. To avoid potential feeding state-related behavioral changes we stopped our tests at 7 dpf, although larvae can survive up to 10 days without food^57^. It would, however, be interesting to investigate whether the observed consistency persists until adulthood, as for adult zebrafish activity levels have also been shown to be consistent over several days^58^, and whether social interactions among the individuals could influence the consistency of this behavior. For some fish species, such as the Amazon molly (*Poecilia Formosa)^42^* or the mangrove killifish (*Kyptolebias marmoratus)^59^* direct social experience did not influence the repeatability of behavior in individuals, despite the importance of social interactions in these species^60^. Yet in other species (e.g. guppy, *Poecilia reticulata* ^61^; rainbow trout, *Onchorhyncus mykiss* ^62^ and cichlid, *Neolamprologus pulcher*^63^), the social environment was shown to affect consistent individual behaviors and the development of animal personalities^64^.

Variability in zebrafish locomotor activity has previously been reported to decrease in the afternoon between 13.00 pm and 15.30 pm ^11^, a result that we could not recapitulate with our data. In fact, during dark intervals we found that the variability was moderately higher in the afternoons at 5 and 7 dpf. A possible explanation for this discrepancy is the difference in protocols as well as light conditions used between our study and the one performed by MacPhail *et al*.. When testing for time of the day effect MacPhail *et al*. kept their larvae under infrared light in constant darkness throughout the period of testing, i.e. from 10.00 am to 15.30 pm, which might have resulted in less variability of the larvae’s locomotor activity when tested in the afternoon. In contrast, in our study, larvae were maintained in normal light conditions between the two test points within one day, in order to mimic natural circadian light conditions and rhythms as closely as possible. Our data also revealed a high intra-individual consistency between morning and afternoon locomotor activity for all days tested and under all light conditions. This within-day consistency allows researchers interested in acute effects to record the baseline activity before the manipulation and thus calculating the effects more precisely.

By 5 dpf, zebrafish larvae perform a repertoire of simple sensorimotor behaviors that operate on characterized and accessible neural circuits^34,65–67^. For example, exposure to abrupt acoustic stimuli elicits a startle response, an evolutionary conserved and stereotyped yet modifiable behavior. Previous research has shown that zebrafish larvae habituate to a startle response. Best *et al*. demonstrated that zebrafish larvae (7 dpf) exhibit frequent reduction in response to a series of acoustic stimuli^68^. Wild-type larvae at 5dpf have also been shown to rapidly reduce their startle response initiation and stereotypically habituate by more than 80% when exposed to a series of acoustic stimuli^69^. In our study, the larvae’s startle response was very inconsistent and unpredictable, for either the dark flashes or tapping stimulus. There was one exception, with the responses of the first tapping stimulus showing moderate correlation between day 6 and 7. This correlation weakened over the 4 stimuli potentially as a result of inter-individual differences in startle response habituation, where some larvae habituated to the stimulus while others did not. Such individuality in habituation was reported for the acoustic startle response by Pantoja *et al*. who showed that the degree of habituation, despite being diverse, is stable and heritable for an individual^70^.

Although our data indicates occasional occurrences of habituation, consistency for an individual, as seen in previous studies, was not apparent. This may be due to the differing startle protocols and different well sizes used for all the studies. The inter-stimulus intervals lasted from 1 second^68,69^ to 5 seconds^70^ in the other studies, while in our study it was 90 seconds, which is much less likely to induce habituation. Moreover, in our setup, the response may have been limited by the size of the well. For example, larvae that are located close to the well edge when the stimulus is triggered, may respond with a small, or large swimming distance depending on the direction of turning. Following the radial index over time indicates that larvae reach the edge of the well during a startle response (see Fig. 1a - 1c). Therefore, intra-individual consistency in startle response habituation might be masked in a standard 48-well plate.

Previous studies have seen strong links between the behavior and morphology of a fish. A study by Hawkins and Quinn (1996) investigated if morphological and physiological traits explained variations in critical swimming speed and found that the best swimming fish had longer caudal regions than the poorer swimmers^71^. Larger brown trout have also been shown to have greater stamina and attained higher swimming speeds than smaller fish, along with maximum swimming speed additionally correlating with fish size^72^. Studies with juvenile zebrafish have shown that individual body size had a strong effect on the activity-boldness relationship, where smaller fish were bolder and less active while larger fish were more cautious and active^73^. In our study, despite strong intra-individual consistency, no such links between behavioural movements and morphological or cardiophysiological parameters were observed under any of the conditions measured. This difference between our data and this literature may be as a result of our study occurring over development. However, other literature is in line with our findings in terms of the lack of this link, as for the bluegill sunfish (*Lepomis macrochirus*), neither boldness nor locomotion activity correlated to the body size or condition of the fish^74^. Importantly, locomotor activity was shown to be independent of weight and body length in adult zebrafish^58^. Therefore, the link between morphology and behaviour may be dependent on the age, conditions and type of behaviour investigated.

Scientific research is constantly under intense scrutiny, specifically for the occurrence of irreproducible and non-comparable findings. In particular, high-throughput behavioral tests frequently result in inconsistent findings, which researchers attribute to poor quality science and non-standardized protocols^9–13^. However, this problem also strongly links to the lack of understanding of the variability of basic behavioral patterns, as fish fundamentally change their swimming behavior over time^1–7,75^. Considering such changes along with the variability would allow the design of a statistically more robust experiment yielding relevant and reliable results. The data from our study is important in helping the development and generation of reproducible zebrafish behavioral data, such as those generated in neurotoxicity or drug discovery tests^8,76–79^. Our results indicate that the behavioral locomotor machinery of an individual, although still under maturation, becomes stable over those key larval stages that are frequently used for testing (6-7 dpf). This stability manifests after the establishment of the beat-and glide swim mode and strengthens when the locomotor network calls upon during darkness-induced exploration, and results in consistent period at 6dpf under dark intervals. In addition, measures between morning and afternoon showed a high intra-individual consistency for all days and light conditions tested. The revealed intra-individual consistency provides some basis to improve the estimation of acute behavioral effects of substances and other types of treatments through pre-post exposure measurements. In addition, this study has highlighted areas where high levels of inter- and/or intra- individual variability occur, specifically for the response to a startle stimulus and morphological and cardiophysiological features of the larvae which should be accounted for when used in future studies. This data not only highlights the need to consider the design and experimental setup/conditions but also provides a basis to allow future studies to account for variability when using zebrafish locomotor behavior. This in turn could help to encourage the inclusion of variability as an additional endpoint, as it might provide new insights into the understanding of an individual’s response to an external challenge.

## Material and Methods

### Zebrafish husbandry

Mixed wildtype (WM) zebrafish (*Danio rerio*), originally obtained from crosses between AB, Tübingen and a pet shop population (OBI, Leipzig, Germany) were maintained under standard conditions^80^ in accordance with the Swiss animal protection law. Adult fish were maintained in a mixed sex Mass Embryo Production System (Aquatic Habitats^®^, Pentair Aquatic Eco-systems, USA), linked to a recirculating flow-through supplied with a mixture of tap and desalted water (1:1) at 26°C ± 1, under a 14:10h light/dark cycle. Adult fish were fed twice daily to satiation from a combination of dry flakes (Tetra, Germany) and live food (*Artemia nauplii*). Group crosses resulted in larvae for the behavioral trials, where the eggs were collected approximately 1 hour post fertilization (hpf). They were rinsed and incubated in aerated artificial freshwater (according to ISO-7346/3 guideline^81^) and unfertilized eggs were removed during the blastula stage as described by Kimmel *et al*. (1995)^82^. Fish were raised in petri dishes, approx. 50 per dish, until needed for behavioral experiments in an incubator with the same light and temperature conditions, as mentioned above, in ISO artificial freshwater, which was changed regularly. All experiments were carried out in accordance with the animal protection guidelines and experiments with larvae were approved by the cantonal veterinary office under the license ZH168/17.

### Behavioral tracking and recording

At 4 dpf, larvae were distributed into 48-well-plates (Greiner Bio-One, Austria), where 1 larva was placed into each well containing 500 μl of fresh ISO water. Fish were moved in the morning and were returned to the housing incubator until the following day when behavioral experiments occurred. Behavior was recorded using the DanioVision Observation Chamber (v. DVOC-0040T; Noldus, Netherlands), which consists of a Gigabit Ethernet video camera, infrared and white light sources, and a transparent multi-well plate holder. The camera output was fed into a standard PC system with the EthoVision XT13 software (version 13.0.1220, Noldus, Netherlands) which created videos to later be analyzed for the fish movement.

Larvae were subjected to different protocols to allow thorough assessment of their movement. The first protocol consisted of an acclimation period of 20 minutes, to allow the fish to adjust to the Noldus set up and allow their baseline movement to settle, followed by a 20 minutes measurement of spontaneous swimming behavior (referred to as “spontaneous” throughout the manuscript). This was then followed by alternating dark and light periods at 10 minutes each (2x dark periods referred to as “dark intervals”, 2x light periods referred to as “light intervals”). Immediately following this protocol, the larvae’s responses to a short pulse of darkness was recorded as follows: Larvae were left for 90 seconds in light before being subjected to a 1 second pulse of darkness, which was repeated 4 times, with an inter-stimulus-interval (ISI) of 90 seconds to allow fish to settle down and reach the baseline in between each stimulus. This ISI was selected as it was shown by Pantoja *et al*. (2016)^70^ to be sufficient to not cause fish to habituate to an acoustic stimulus. The same pattern was used for the tapping protocol but this time the stimulus was produced using the inbuilt DanioVision Tapping Device (Noldus, Netherlands)^38^ which produces an acoustic vibrational stimulus. The swimming protocol was recorded at 30 frames per second while the startle protocols were recorded at 60 frames per second. All of the behavior protocols were run at 9am and 2pm, on 5, 6 and 7 dpf, with all the protocols being the same for all measurements carried out. Between the measurements at 9am and 2pm, the well-plate was maintained under light conditions in the testing room to most closely mimic their normal diurnal cycle. The 3 days experiment was repeated 3 times at different periods, producing a sample size of n = 132 larvae for behavioral analysis. There was no effect of the rep on the behavior of the larvae for any of the conditions tested (spontaneous: χ^2^ (1) = 0.18, p = 0.672; dark intervals: χ^2^ (1) = 0.07, p = 0.787; spontaneous: χ^2^ (1) = 0.83, p = 0.361). In addition, there was no effect of location of the fish within the plate for any of the conditions tested (e.g. spontaneous swimming-well: χ^2^ (47) = 44.82, p = 0.563; column: χ^2^ (1) = 0.16, p = 0.693; row: χ^2^ (5) = 4.79, p = 0.442).

### Heart rate and morphological measurements

At 4pm on each of the 3 days, videos were recorded of each larva for the measurement of heart rates and morphometric parameters. Larvae were anesthetized with 160 mg/L of ethyl 3-aminobenzoate methanesulfonate (MS222; Sigma-Aldrich). A 15 seconds video of each larva in the lateral position was then captured, at 30 frames per second, using a Basler acA2000-165μM camera mounted on a Leica S8APO stereo microscope. Videos were recorded using the Media Recorder 4 software (version 4.0; Noldus, Netherlands). After a suitable video had been taken, larvae were immediately moved into a bath of 100% air saturated ISO water to allow recovery from the anesthetic. Recovered larvae were then moved to the same position of a new 48-well-plate containing 500 μl of ISO water in each well and placed back into the incubator until the next behavioral measurements were taken.

From the videos collected, the heart rate and morphology parameters were measured using DanioScope Software (version 1.2.206; Noldus, Netherlands). After manually selecting the heart area in the video, the software calculated the number of beats per minute using a power plot spectrum of the frequencies extracted from an activity signal. For each larva, 3 heart rate measurements were taken for each day from the same video and the average of these was taken as the heart rate for that larvae on that day. The morphology parameters measured were body length (from nose to tip of the tail; mm) and swim bladder (size of the swim bladder from the lateral view of the larvae; mm^2^), using the DanioScope software. The same calibration profile was used for all images, to ensure comparability between each image.

### Data and statistical analysis

Tracking of the fish by the EthoVision software was carried out from the videos in a non-live tracking mode, allowing a static subtraction of the background and reducing tracking artifacts. The same settings were ensured to give consistency across the different day, times and replicates. To characterize the swimming behaviors during the spontaneous, dark and light intervals, the activity and radial index were calculated. The activity index represents the percentage of movement by each larva within one-second intervals and was calculated within the software program using 2.00 to 1.75 cm/s as a threshold. The radial index indicated where the larva moved within the well and was calculated using the distance from center of the well (calculated in the EthoVision software) and dividing it by the radius of 5.725 mm. To calculate the intra-individual coefficient of variance, standard deviation was divided by the mean activity for each individual larva.

For the startle response strength, the distance moved per second was used instead of the activity index. This did not affect the significance or trends that were observed with the activity index but allowed a more detailed view of the specific movements of the fish. The period one-second post stimulus was taken to analyze the distance moved by the fish while performing a startle response. To calculate the strength of each response with respect to spontaneous movement, a baseline for each fish was determined before each stimulus. The distance moved per second during a 40 second period, 10 seconds before the startle stimulus, was averaged and SD added to give baseline level of distance moved before the stimulus. The response strength was then calculated by subtracting the baseline from the distance moved during the startle stimulus. Using this baseline, the habituation index was calculated from ratios between the first strength response and either the second, third or fourth strength response. The sum of all ratios was taken as the habituation score for that larva. This score was calculated for each individual for each day and time.

All statistical analysis was carried out in ‘RStudio’ (version 1.1.453, USA). To carry out the correlation analysis, the Pearson’s correlation coefficient, r, was calculated and reported, and the p-values to test this correlation obtained from a two-sided t-test. A linear regression model was used to draw lines of best fit for the scatter plots. To test between group differences and check for differences between conditions, analysis of variance models were carried out using the minimum adequate model approach, where model simplification using F test occurred based on analysis of deviance. Linear mixed effect analysis was carried out to test for positional effects of the well, as well as effects from the different repeats. Well location, column, row, edge and repeat were entered as fixed effects, while individual larva were entered as random effects. P values were obtained by likelihood ratio tests of the full model with the effect in question against the model without the effect. For all tests, data were considered statistically significant when p < 0.05.

## Data Availability

The datasets generated and analyzed during the current study are available in the ERIC repository, https://data.eawag.ch/.

## Acknowledgements

We would like to thank Pascal Reichlin, Marianne Zimmermann and Hidir Sengül for expert fish care. We would also like to thank Roman Li and Anze Zupanic for critical comments on the manuscript. We are grateful to the whole Utox department for fruitful discussions and helpful suggestions throughout the study.

## Author Contributions

C.M.vB conceived the study; K.T.K and C.M.vB designed and carried out the preliminary experiments; J.A.F performed the experiments and carried out the analysis; C.P.Z and J.A.F programmed the analysis tools; J.A.F and C.M.vB wrote the manuscript. All authors critically revised the manuscript and confirmed the last version.

## Additional Information

The authors declare no competing interests.

## References

1. Naganawa, Y. & Hirata, H. Developmental transition of touch response from slow muscle-mediated coilings to fast muscle-mediated burst swimming in zebrafish. Dev Biol 355, 194–204, doi:10.1016/j.ydbio.2011.04.027 (2011).

2. Saint-Amant, L. & Drapeau, P. Time course of the development of motor behaviors in the zebrafish embryo. J Neurobiol 37, 622–632 (1998).

3. Budick, S. A. & O’Malley, D. M. Locomotor repertoire of the larval zebrafish: swimming, turning and prey capture. J Exp Biol 203, 2565–2579 (2000).

4. Buss, R. R. & Drapeau, P. Synaptic drive to motoneurons during fictive swimming in the developing zebrafish. J Neurophysiol 86, 197–210, doi:10.1152/jn.2001.86.1.197 (2001).

5. Brustein, E. et al. Steps during the development of the zebrafish locomotor network. J Physiol Paris 97, 77–86, doi:10.1016/j.jphysparis.2003.10.009 (2003).

6. Drapeau, P. et al. Development of the locomotor network in zebrafish. Prog Neurobiol 68, 85–111 (2002).

7. Fero, K., Yokogawa, T. & Burgess, H. A. The Behavioral Repertoire of Larval Zebrafish. Zebrafish Models in Neurobehavioral Research, 249–291, doi:10.1007/978-1-60761-922-2_12 (2011).

8. Farrell, T. C. et al. Evaluation of spontaneous propulsive movement as a screening tool to detect rescue of Parkinsonism phenotypes in zebrafish models. Neurobiol Dis 44, 9–18, doi:10.1016/j.nbd.2011.05.016 (2011).

9. Ingebretson, J. J. & Masino, M. A. Quantification of locomotor activity in larval zebrafish: considerations for the design of high-throughput behavioral studies. Front Neural Circuits 7, 109, doi:10.3389/fncir.2013.00109 (2013).

10. Legradi, J., el Abdellaoui, N., van Pomeren, M. & Legler, J. Comparability of behavioural assays using zebrafish larvae to assess neurotoxicity. Environ Sci Pollut Res Int 22, 16277–16289, doi:10.1007/s11356-014-3805-8 (2015).

11. MacPhail, R. C. et al. Locomotion in larval zebrafish: Influence of time of day, lighting and ethanol. Neurotoxicology 30, 52–58, doi:10.1016/j.neuro.2008.09.011 (2009).

12. Melvin, S. D., Petit, M. A., Duvignacq, M. C. & Sumpter, J. P. Towards improved behavioural testing in aquatic toxicology: Acclimation and observation times are important factors when designing behavioural tests with fish. Chemosphere 180, 430–436, doi:10.1016/j.chemosphere.2017.04.058 (2017).

13. Melvin, S. D. & Wilson, S. P. The utility of behavioral studies for aquatic toxicology testing: a meta-analysis. Chemosphere 93, 2217–2223, doi:10.1016/j.chemosphere.2013.07.036 (2013).

14. Padilla, S., Hunter, D. L., Padnos, B., Frady, S. & MacPhail, R. C. Assessing locomotor activity in larval zebrafish: Influence of extrinsic and intrinsic variables. Neurotoxicol Teratol 33, 624–630, doi:10.1016/j.ntt.2011.08.005 (2011).

15. John, O. P., Robins, R. & Pervin, L. A. Handbook of Personality. Theory and Research. (1999).

16. Larsen, J. R. & Buss, D. Personality Psychology: Domains of Knowledge about Human Nature. (2005).

17. Drent, P. J., van Oers, K. & van Noordwijk, A. J. Realized heritability of personalities in the great tit (Parus major). Proc Biol Sci 270, 45–51, doi:10.1098/rspb.2002.2168 (2003).

18. Groothuis, T. G. & Carere, C. Avian personalities: characterization and epigenesis. Neurosci Biobehav Rev 29, 137–150, doi:10.1016/j.neubiorev.2004.06.010 (2005).

19. Conrad, J. L., Weinersmith, K. L., Brodin, T., Saltz, J. B. & Sih, A. Behavioural syndromes in fishes: a review with implications for ecology and fisheries management. J Fish Biol 78, 395–435, doi:10.1111/j.1095-8649.2010.02874.x (2011).

20. Gosling, S. D. From mice to men: what can we learn about personality from animal research? Psychol Bull 127, 45–86 (2001).

21. Wolf, M. & Weissing, F. J. Animal personalities: consequences for ecology and evolution. Trends in Ecology & Evolution 27, 452–461, doi:10.1016/j.tree.2012.05.001 (2012).

22. Kappeler, P. & Kraus, C. Levels and mechanisms of behavioural variability. Animal Behaviour: Evolution and Mechanisms, 655–684, doi:10.1007/978-3-642-02624-9_21 (2010).

23. Laskowski, K. L. & Bell, A. M. Strong personalities, not social niches, drive individual differences in social behaviours in sticklebacks. Anim Behav 90, 287–295, doi:10.1016/j.anbehav.2014.02.010 (2014).

24. Nikinmaa, M. & Anttila, K. Individual variation in aquatic toxicology: Not only unwanted noise. Aquat Toxicol 207, 29–33, doi:10.1016/j.aquatox.2018.11.021 (2019).

25. Dingemanse, N. J., Kazem, A. J., Reale, D. & Wright, J. Behavioural reaction norms: animal personality meets individual plasticity. Trends Ecol Evol 25, 81–89, doi:10.1016/j.tree.2009.07.013 (2010).

26. Wolf, M. & Weissing, F. J. An explanatory framework for adaptive personality differences. Philos Trans R Soc Lond B Biol Sci 365, 3959–3968, doi:10.1098/rstb.2010.0215 (2010).

27. Bennett, A. F. Interindividual variability: an underutilized resource. New Directions in Ecological Physiology 19, 147–169 (1987).

28. Sih, A., Bell, A. M., Johnson, J. C. & Ziemba, R. E. Behavioral syndromes: an intergrative overiew. Q Rev Biol 79, 241–277 (2004).

29. Roche, D. G., Careau, V. & Binning, S. A. Demystifying animal ‘personality’ (or not): why individual variation matters to experimental biologists. J Exp Biol 219, 3832–3843, doi:10.1242/jeb.146712 (2016).

30. Williams, T. D. Individual variation in endocrine systems: moving beyond the ‘tyranny of the Golden Mean’. Philos Trans R Soc Lond B Biol Sci 363, 1687–1698, doi:10.1098/rstb.2007.0003 (2008).

31. Piersma, T. & Drent, J. P. Phenotypic flexibility and the evolution of organismal design. Trends in Ecology & Evolution 18, 228–233, doi:10.1016/S0169-5347(03)00036-3 (2003).

32. Stamps, J. A., Briffa, M. & Biro, P. A. Unpredictable animals: individual differences in intra-individual variability (IIV). Animal Behaviour 83, 1325–1334, doi:10.1016/j.anbehav.2012.02.017 (2012).

33. Hayes, J. P. & Jenkins, S. H. Individual Variation in Mammals. Journal of Mammalogy 78, 274–293, doi:10.2307/1382882 (1997).

34. Burgess, H. A. & Granato, M. Modulation of locomotor activity in larval zebrafish during light adaptation. J Exp Biol 210, 2526–2539, doi:10.1242/jeb.003939 (2007).

35. Emran, F., Rihel, J. & Dowling, J. E. A behavioral assay to measure responsiveness of zebrafish to changes in light intensities. J Vis Exp, doi:10.3791/923 (2008).

36. Fernandes, A. M. et al. Deep brain photoreceptors control light-seeking behavior in zebrafish larvae. Curr Biol 22, 2042–2047, doi:10.1016/j.cub.2012.08.016 (2012).

37. Horstick, E. J., Bayleyen, Y., Sinclair, J. L. & Burgess, H. A. Search strategy is regulated by somatostatin signaling and deep brain photoreceptors in zebrafish. BMC Biol 15, 4, doi:10.1186/s12915-016-0346-2 (2017).

38. Noldus, L. P. https://www.noldus.com/daniovision/tapping-device.

39. Roman, A. C. et al. Histone H4 acetylation regulates behavioral inter-individual variability in zebrafish. Genome Biol 19, 55, doi:10.1186/s13059-018-1428-y (2018).

40. Parichy, D. M., Elizondo, M. R., Mills, M. G., Gordon, T. N. & Engeszer, R. E. Normal table of postembryonic zebrafish development: staging by externally visible anatomy of the living fish. Dev Dyn 238, 2975–3015, doi:10.1002/dvdy.22113 (2009).

41. Robertson, G. N., McGee, C. A., Dumbarton, T. C., Croll, R. P. & Smith, F. M. Development of the swimbladder and its innervation in the zebrafish, Danio rerio. J Morphol 268, 967–985, doi:10.1002/jmor.10558 (2007).

42. Bierbach, D., Laskowski, K. L. & Wolf, M. Behavioural individuality in clonal fish arises despite near-identical rearing conditions. Nat Commun 8, 15361, doi:10.1038/ncomms15361 (2017).

43. Freund, J. et al. Emergence of individuality in genetically identical mice. Science 340, 756–759, doi:10.1126/science.1235294 (2013).

44. Baker, M. R., Goodman, A. C., Santo, J. B. & Wong, R. Y. Repeatability and reliability of exploratory behavior in proactive and reactive zebrafish, Danio rerio. Sci Rep 8, 12114, doi:10.1038/s41598-018-30630-3 (2018).

45. de Esch, C. et al. Locomotor activity assay in zebrafish larvae: influence of age, strain and ethanol. Neurotoxicol Teratol 34, 425–433, doi:10.1016/j.ntt.2012.03.002 (2012).

46. Gao, Y. et al. Computational classification of different wild-type zebrafish strains based on their variation in light-induced locomotor response. Comput Biol Med 69, 1–9, doi:10.1016/j.compbiomed.2015.11.012 (2016).

47. Lange, M. et al. Inter-individual and inter-strain variations in zebrafish locomotor ontogeny. PLoS One 8, e70172, doi:10.1371/journal.pone.0070172 (2013).

48. Moretz, J. A., Martins, E. P. & Robison, B. D. Behavioral syndromes and the evolution of correlated behavior in zebrafish. Behavioral Ecology 18, 556–562, doi:10.1093/beheco/arm011 (2007).

49. Vignet, C. et al. Systematic screening of behavioral responses in two zebrafish strains. Zebrafish 10, 365–375, doi:10.1089/zeb.2013.0871 (2013).

50. Dunn, T. W. et al. Brain-wide mapping of neural activity controlling zebrafish exploratory locomotion. Elife 5, e12741, doi:10.7554/eLife.12741 (2016).

51. Gahtan, E., Tanger, P. & Baier, H. Visual prey capture in larval zebrafish is controlled by identified reticulospinal neurons downstream of the tectum. J Neurosci 25, 9294–9303, doi:10.1523/JNEUROSCI.2678-05.2005 (2005).

52. Preuss, S. J., Trivedi, C. A., vom Berg-Maurer, C. M., Ryu, S. & Bollmann, J. H. Classification of object size in retinotectal microcircuits. Curr Biol 24, 2376–2385, doi:10.1016/j.cub.2014.09.012 (2014).

53. Burgess, H. A., Schoch, H. & Granato, M. Distinct retinal pathways drive spatial orientation behaviors in zebrafish navigation. Curr Biol 20, 381–386, doi:10.1016/j.cub.2010.01.022 (2010).

54. Mueller, K. P. & Neuhauss, S. C. Behavioral neurobiology: how larval fish orient towards the light. Curr Biol 20, R159–161, doi:10.1016/j.cub.2009.12.028 (2010).

55. Clift, D., Richendrfer, H., Thorn, R. J., Colwill, R. M. & Creton, R. High-throughput analysis of behavior in zebrafish larvae: effects of feeding. Zebrafish 11, 455–461, doi:10.1089/zeb.2014.0989 (2014).

56. Dametto, F. S. et al. Feeding regimen modulates zebrafish behavior. PeerJ 6, e5343, doi:10.7717/peerj.5343 (2018).

57. Hernandez, R. E., Galitan, L., Cameron, J., Goodwin, N. & Ramakrishnan, L. Delay of Initial Feeding of Zebrafish Larvae Until 8 Days Postfertilization Has No Impact on Survival or Growth Through the Juvenile Stage. Zebrafish 15, 515–518, doi:10.1089/zeb.2018.1579 (2018).

58. Tran, S. & Gerlai, R. Individual differences in activity levels in zebrafish (Danio rerio). Behav Brain Res 257, 224–229, doi:10.1016/j.bbr.2013.09.040 (2013).

59. Edenbrow, M. & Croft, D. P. Environmental and genetic effects shape the development of personality traits in the mangrove killifish Kryptolebias marmoratus. Oikos 122, 667–681, doi:10.1111/j.1600-0706.2012.20556.x (2013).

60. Schlupp, I. Chapter 5 Behavior of Fishes in the Sexual/Unisexual Mating System of the Amazon Molly (Poecilia formosa). Advances in the Study of Behavior 39, 153–183, doi:10.1016/S0065-3454(09)39005-1 (2009).

61. Chapman, B. B., Ward, A. J. W. & Krause, J. Schooling and learning: early social environment predicts social learning ability in the guppy, Poecilia reticulata. Animal Behaviour 76, 923–929, doi:10.1016/j.anbehav.2008.03.022 (2008).

62. Frost, A. J., Winrow-Giffen, A., Ashley, P. J. & Sneddon, L. U. Plasticity in animal personality traits: does prior experience alter the degree of boldness? Proc Biol Sci 274, 333–339, doi:10.1098/rspb.2006.3751 (2007).

63. Arnold, C. & Taborsky, B. Social experience in early ontogeny has lasting effects on social skills in cooperatively breeding cichlids. Animal Behaviour 79, 621–630, doi:10.1016/j.anbehav.2009.12.008 (2010).

64. Bergmuller, R. & Taborsky, M. Animal personality due to social niche specialisation. Trends Ecol Evol 25, 504–511, doi:10.1016/j.tree.2010.06.012 (2010).

65. Eaton, R. C., Bombardieri, R. A. & Meyer, D. L. The Mauthner-initiated startle response in teleost fish. J Exp Biol 66, 65–81 (1977).

66. Kimmel, C. B., Patterson, J. & Kimmel, R. O. The development and behavioral characteristics of the startle response in the zebra fish. Dev Psychobiol 7, 47–60, doi:10.1002/dev.420070109 (1974).

67. Troconis, E. L. et al. Intensity-dependent timing and precision of startle response latency in larval zebrafish. J Physiol 595, 265–282, doi:10.1113/JP272466 (2017).

68. Best, J. D. et al. Non-associative learning in larval zebrafish. Neuropsychopharmacology 33, 1206–1215, doi:10.1038/sj.npp.1301489 (2008).

69. Wolman, M. A., Jain, R. A., Liss, L. & Granato, M. Chemical modulation of memory formation in larval zebrafish. Proc Natl Acad Sci U S A 108, 15468–15473, doi:10.1073/pnas.1107156108 (2011).

70. Pantoja, C. et al. Neuromodulatory Regulation of Behavioral Individuality in Zebrafish. Neuron 91, 587–601, doi:10.1016/j.neuron.2016.06.016 (2016).

71. Hawkins, D. K. & Quinn, T. P. Critical swimming velocity and associated morphology of juvenile coastal cutthroat trout (Oncorhynchus clarki clarki), steelhead trout (Oncorhynchus mykiss), and their hybrids. Canadian Journal of Fisheries and Aquatic Sciences 53, 1487–1496, doi:10.1139/f96-085 (1996).

72. Ojanguren, A. F. & Braña, F. Effects of size and morphology on swimming performance in juvenile brown trout (Salmo trutta L.). Ecology of Freshwater Fish 12, 241–246, doi:10.1046/j.1600-0633.2003.00016.x (2003).

73. Roy, T. & Bhat, A. Population, sex and body size: determinants of behavioural variations and behavioural correlations among wild zebrafish Danio rerio. R Soc Open Sci 5, 170978, doi:10.1098/rsos.170978 (2018).

74. Wilson, A. D. M. & Godin, J.-G. J. Boldness and intermittent locomotion in the bluegill sunfish, Lepomis macrochirus. Behavioral Ecology 21, 57–62, doi:10.1093/beheco/arp157 (2009).

75. Colwill, R. M. & Creton, R. Locomotor behaviors in zebrafish (Danio rerio) larvae. Behav Processes 86, 222–229, doi:10.1016/j.beproc.2010.12.003 (2011).

76. Ali, S., Champagne, D. L. & Richardson, M. K. Behavioral profiling of zebrafish embryos exposed to a panel of 60 water-soluble compounds. Behav Brain Res 228, 272–283, doi:10.1016/j.bbr.2011.11.020 (2012).

77. de Esch, C., Slieker, R., Wolterbeek, A., Woutersen, R. & de Groot, D. Zebrafish as potential model for developmental neurotoxicity testing: a mini review. Neurotoxicol Teratol 34, 545–553, doi:10.1016/j.ntt.2012.08.006 (2012).

78. Legradi, J. B. et al. An ecotoxicological view on neurotoxicity assessment. Environmental Sciences Europe 30, 46, doi:10.1186/s12302-018-0173-x (2018).

79. Tierney, K. B. Behavioural assessments of neurotoxic effects and neurodegeneration in zebrafish. Biochim Biophys Acta 1812, 381–389, doi:10.1016/j.bbadis.2010.10.011 (2011).

80. Westerfield, M. The Zebrafish Book. A Guide for The Laboratory Use of Zebrafish (Danio rerio). Vol. 385 (2000).

81. ISO. Water quality — Determination of the acute lethal toxicity of substances to a freshwater fish [Brachydanio rerio Hamilton-Buchanan (Teleostei, Cyprinidae)] — Part 3: Flow-through method: ISO 7346-3:1996(en). ISO International Standards, 11 (1996).

82. Kimmel, C. B., Ballard, W. W., Kimmel, S. R., Ullmann, B. & Schilling, T. F. Stages of embryonic development of the zebrafish. Dev Dyn 203, 253–310, doi:10.1002/aja.1002030302 (1995).

