## Supplementary Information for "Emergence of consistent intra-individual locomotor patterns during zebrafish development"

**Pages**

**FIGURES**

Supp. Fig. 1 ........ S2

Supp. Fig. 2 ........ S3

Supp. Fig. 3 ........ S4

Supp. Fig. 4 ........ S5

Supp. Fig. 5 ........ S6

**TABLES**

Supp. Tab. 1 ........ S7

Supp. Tab. 2 ........ S8

Supp. Tab. 3 ........ S9

Supp. Tab. 4 ........ S10

Supp. Tab. 5 ........ S11

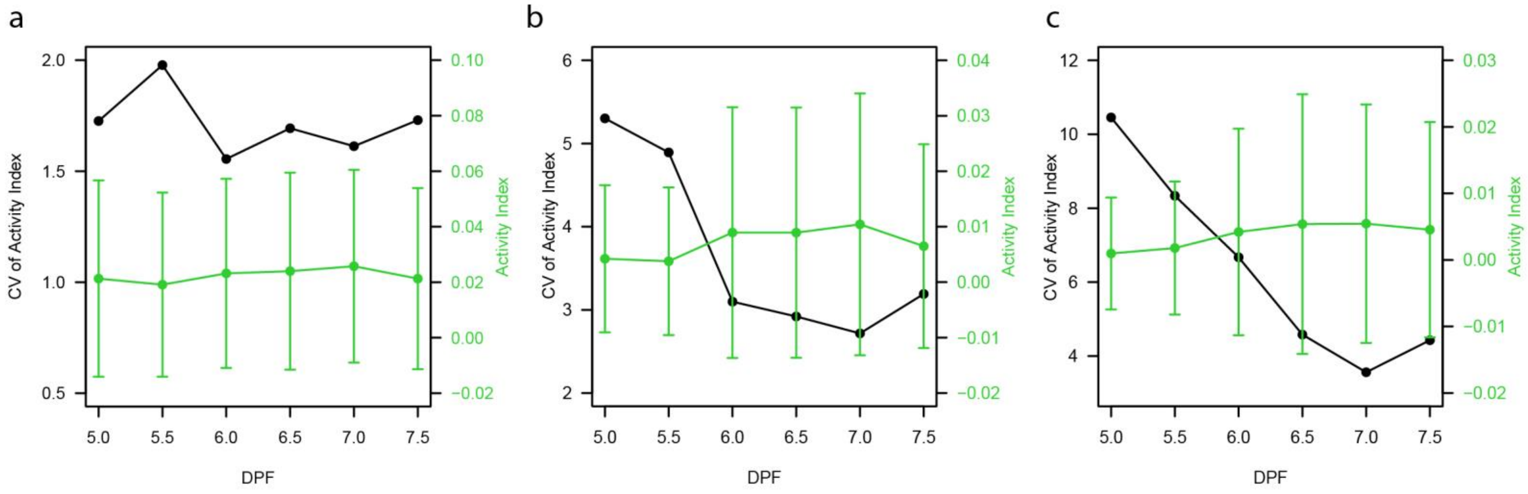


**Supp. Fig 1**: Plots showing the median coefficient of variance (CV) (black) for each individual larva (n = 132) and the average activity index ± standard deviation (green) over the different days and daytimes (days post fertilization), for (a) dark intervals, (b) spontaneous and (c) light intervals.

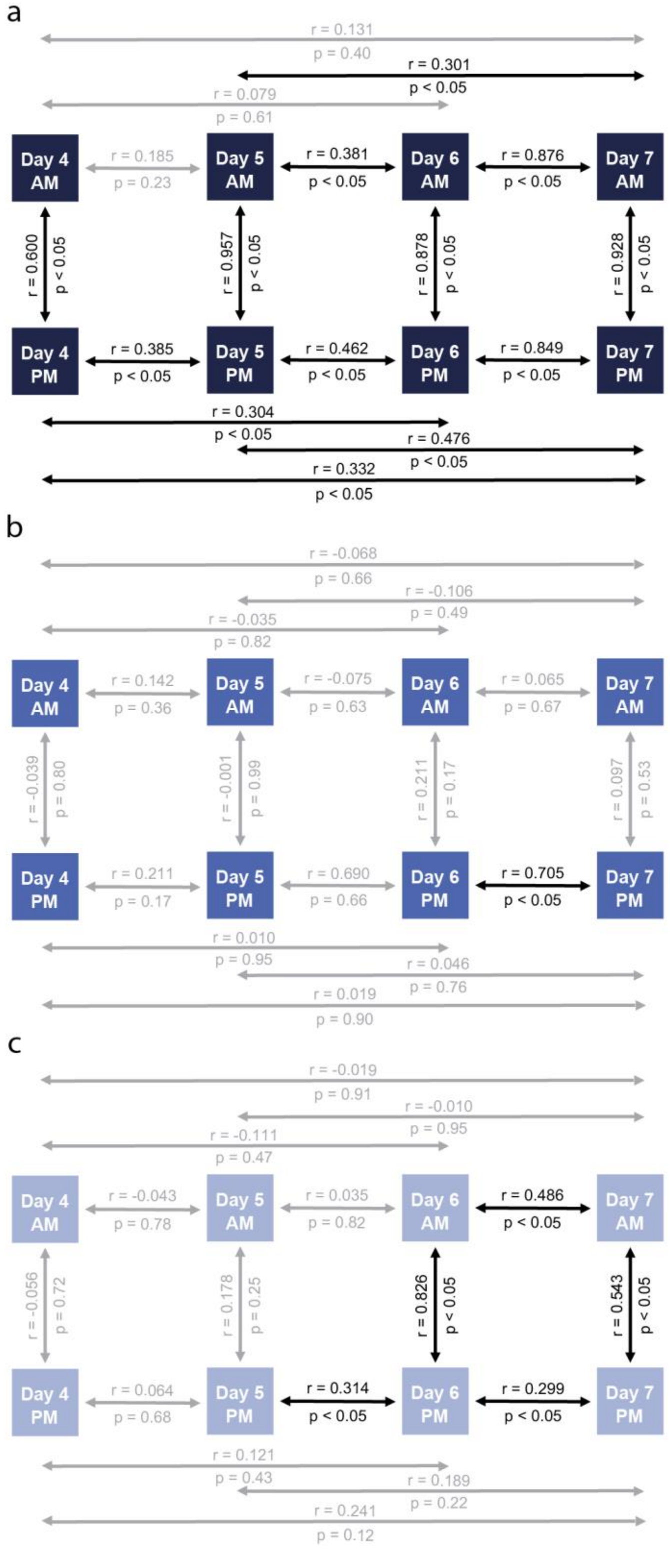


**Supp. Fig 2**: Schematics representing the correlations of the activity index for the 4 day study between different days and time of days for each of the conditions studied, (a) dark intervals, (b) spontaneous and (c) light intervals. All correlation coefficients derived from Pearson's correlations with respective p values.

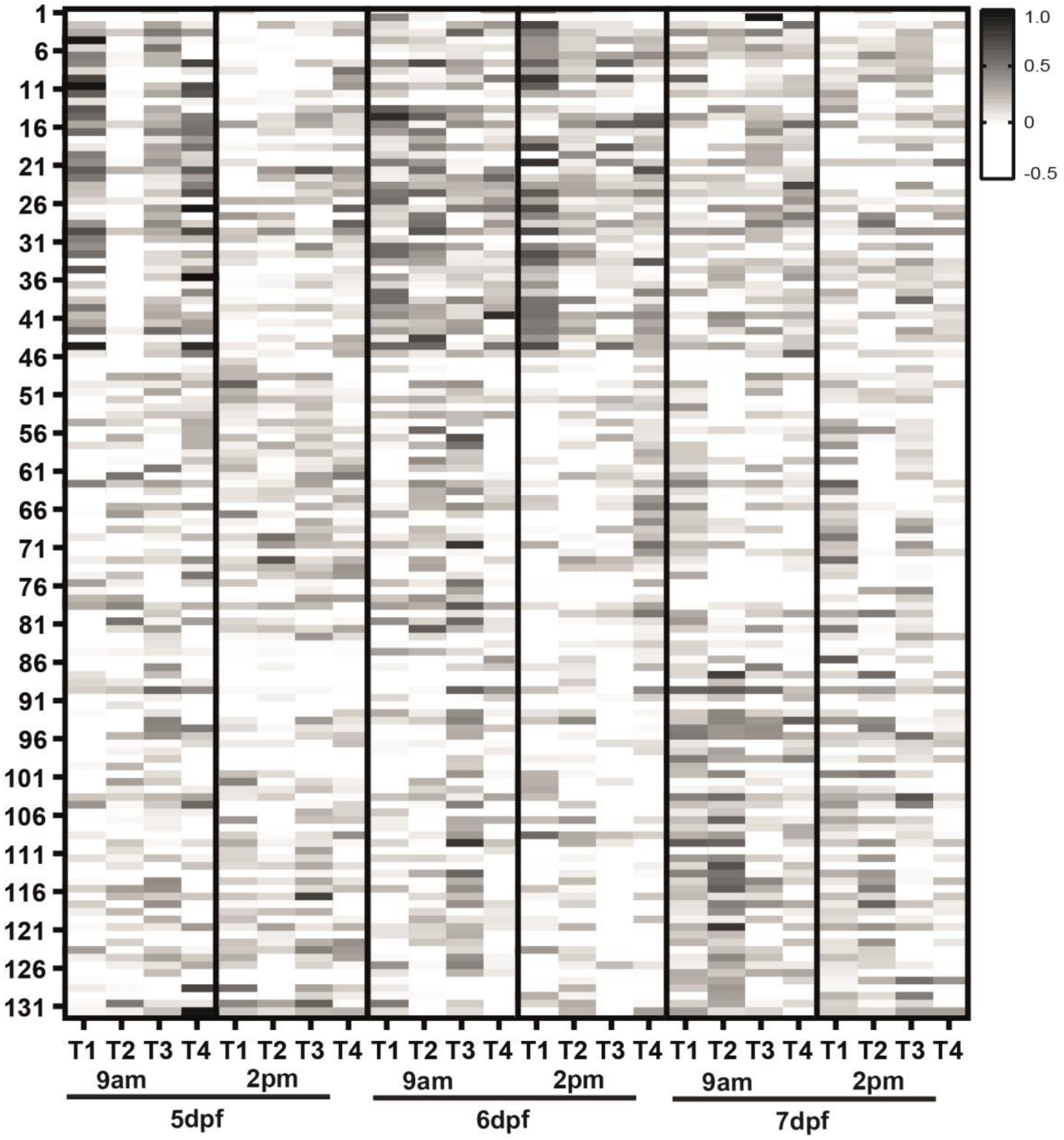


**Supp. Fig. 3:** Heat map representing the change in distance moved with respect to the baseline of each individual larvae for all time points and days measured in response to each of the dark flash stimulus (S1-S4). White represents no response to the stimulus, with the grey scale darkening in a linear scale depending on the strength of the response.

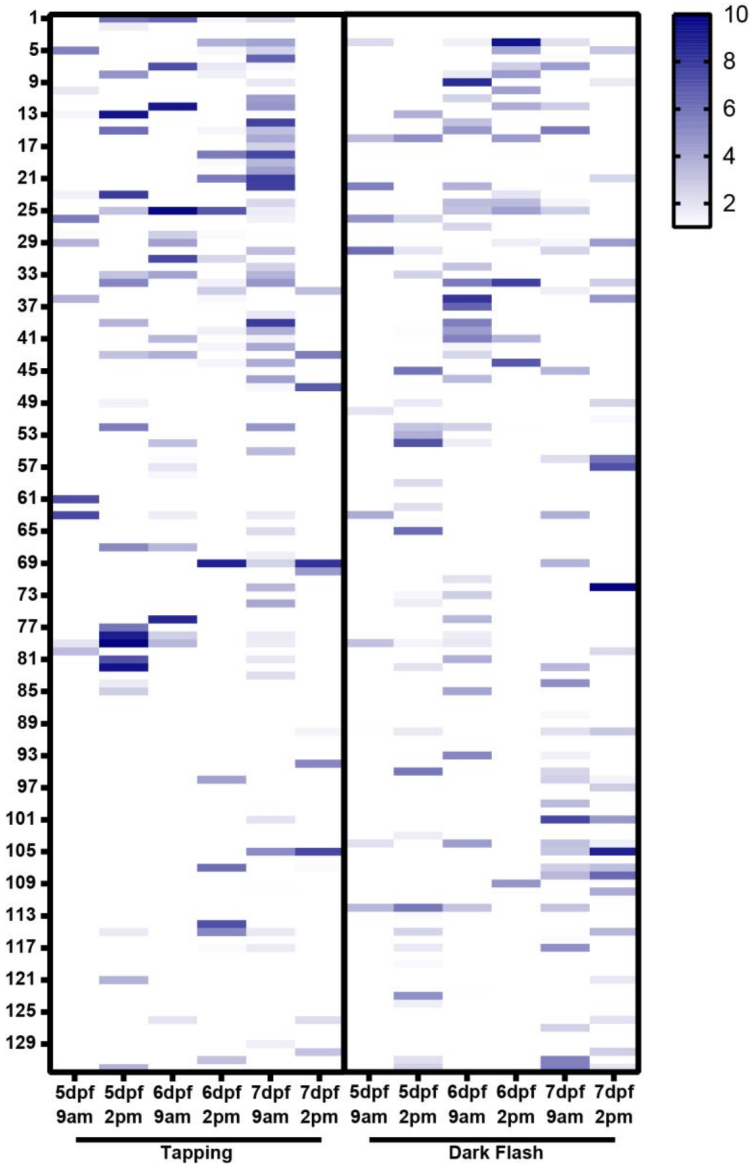


**Supp. Fig. 4:** Heat map showing the habituation index of the 132 individual larvae for all time points and days measured in response to the tapping (left) and dark flash (right) stimuli. White represents no habituation, with blue darkening in a linear scale depending on the strength of habituation calculated.

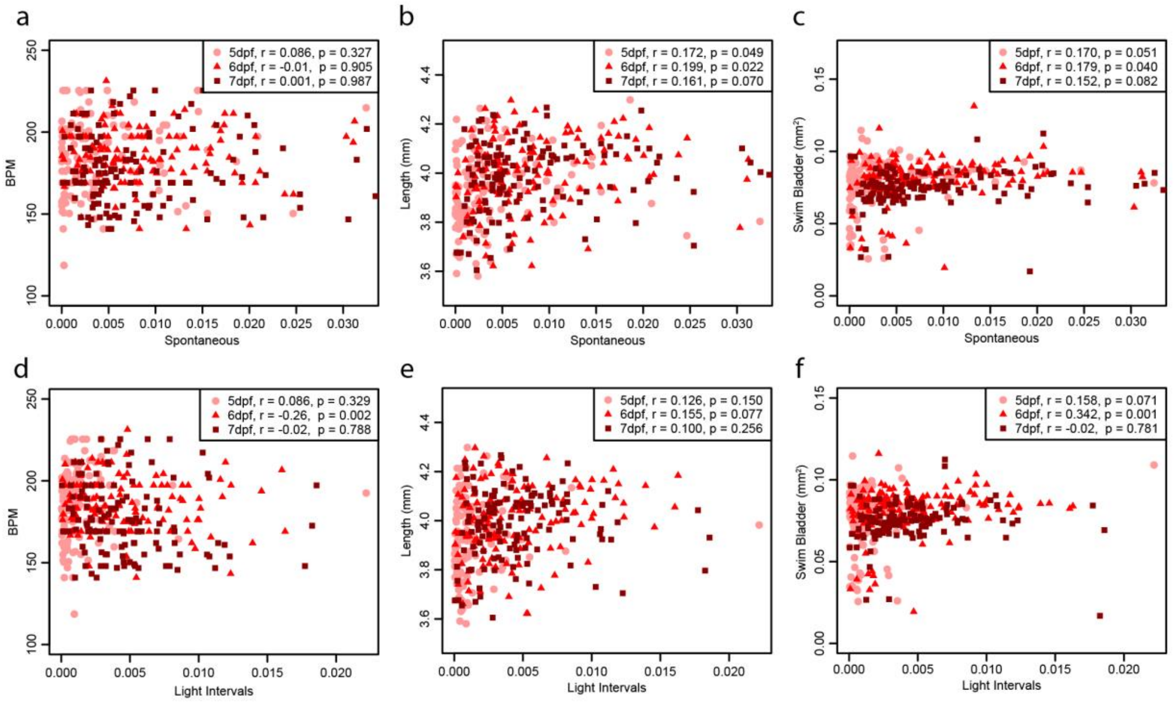


**Supp. Fig. 5:** Comparison of the individual larvae's (a, d) heart rate (in beats per minute, BPM), (b, e) body length (in mm) and (c, f) size of swim bladder (in $mm^2$) to their respective average activity during spontaneous (a, b, c) and light intervals (d, e, f), with each day plotted on each plot (5 days post fertilization (dpf): circle, 6 dpf; triangle and 7 dpf; square). Statistics on the plots represent the Pearson's correlation coefficient and respective p value.

**Supp. Tab. 1:** Table of correlations between the activity and radial index for all conditions tested (spontaneous, dark intervals and light intervals) for all days and times tested. All correlation coefficients were derived from Pearson's correlations, with color scale based on the r value (blue positive correlation and yellow negative correlation). Significant correlations are denoted by the bold font when $p < 0.05$.

| | Day | Time | Activity v Radial | |
|---|---|---|---|---|
| | | | r | p value |
| **Spontaneous** | 5 | 9am | 0.14 | 0.116 |
| | | 2pm | -0.02 | 0.846 |
| | 6 | 9am | **0.21** | **0.014** |
| | | 2pm | **0.24** | **0.005** |
| | 7 | 9am | **0.31** | **$3.54 \times 10^{-4}$** |
| | | 2pm | **0.23** | **0.009** |
| **Dark Intervals** | 5 | 9am | **0.37** | **$1.22 \times 10^{-5}$** |
| | | 2pm | **0.35** | **$3.65 \times 10^{-5}$** |
| | 6 | 9am | **0.34** | **$5.59 \times 10^{-5}$** |
| | | 2pm | **0.32** | **$1.82 \times 10^{-4}$** |
| | 7 | 9am | **0.21** | **0.042** |
| | | 2pm | **0.21** | **0.018** |
| **Light Intervals** | 5 | 9am | 0.04 | 0.638 |
| | | 2pm | 0.08 | 0.350 |
| | 6 | 9am | 0.17 | 0.057 |
| | | 2pm | 0.09 | 0.305 |
| | 7 | 9am | -0.01 | 0.894 |
| | | 2pm | 0.06 | 0.480 |

**Supp. Tab. 2:** Table of correlations between the startle responses (tapping, dark flash, onset and offset) for all days and times tested. Values given are the r values from Pearson's correlations, with color scale based on that r value (blue positive correlation and yellow negative correlation). Significant correlations are denoted by the bold font when $p < 0.05$.

| | | | Tapping | | | | | | Dark Flash | | | | | | Offset | | | | | | Onset | | | | | |
|---|---|---|---|---|---|---|---|---|---|---|---|---|---|---|---|---|---|---|---|---|---|---|---|---|---|---|
| | | | 5 | | 6 | | 7 | | 5 | | 6 | | 7 | | 5 | | 6 | | 7 | | 5 | | 6 | | 7 | |
| | | | 9am | 2pm | 9am | 2pm | 9am | 2pm | 9am | 2pm | 9am | 2pm | 9am | 2pm | 9am | 2pm | 9am | 2pm | 9am | 2pm | 9am | 2pm | 9am | 2pm | 9am | 2pm |
| Tapping | 5 | 9am | | 0.127 | **0.263** | -0.080 | 0.106 | 0.005 | **0.338** | -0.105 | 0.134 | 0.106 | -0.083 | -0.050 | 0.131 | 0.155 | 0.079 | -0.032 | 0.108 | -0.053 | 0.060 | 0.046 | 0.151 | 0.060 | 0.126 | 0.006 |
| | | 2pm | | | **0.395** | 0.092 | **0.230** | 0.044 | **0.188** | 0.060 | **0.216** | 0.166 | -0.275 | -0.025 | -0.033 | -0.008 | 0.126 | -0.076 | **0.172** | -0.026 | -0.119 | -0.041 | -0.027 | -0.064 | **0.172** | -0.007 |
| | 6 | 9am | | | | **0.251** | **0.423** | -0.031 | **0.278** | -0.082 | **0.437** | **0.293** | **-0.242** | -0.103 | 0.048 | -0.073 | 0.085 | -0.157 | 0.012 | -0.160 | -0.074 | -0.037 | 0.140 | -0.027 | **0.245** | 0.025 |
| | | 2pm | | | | | **0.270** | **-0.404** | 0.144 | -0.112 | **0.318** | **0.276** | -0.027 | -0.105 | -0.131 | -0.018 | -0.004 | 0.093 | 0.093 | 0.043 | -0.054 | -0.085 | 0.031 | 0.012 | 0.142 | 0.152 |
| | 7 | 9am | | | | | | -0.106 | **0.482** | -0.103 | **0.486** | **0.563** | **-0.226** | -0.111 | 0.022 | 0.080 | 0.091 | -0.122 | 0.062 | -0.005 | -0.046 | 0.120 | 0.122 | 0.050 | **0.223** | 0.178 |
| | | 2pm | | | | | | | **-0.268** | 0.091 | -0.145 | **-0.320** | 0.018 | **0.266** | 0.130 | 0.061 | 0.092 | -0.150 | 0.092 | 0.035 | -0.130 | 0.064 | 0.135 | -0.081 | -0.070 | **-0.207** |
| Dark Flash | 5 | 9am | | | | | | | | -0.098 | **0.435** | **0.475** | **-0.456** | -0.056 | **0.314** | **0.214** | 0.097 | -0.022 | 0.096 | 0.087 | 0.088 | 0.143 | 0.145 | 0.084 | **0.202** | 0.162 |
| | | 2pm | | | | | | | | | -0.076 | 0.022 | 0.051 | -0.053 | 0.132 | 0.129 | 0.153 | 0.139 | 0.063 | 0.046 | -0.111 | -0.094 | 0.012 | 0.072 | -0.069 | 0.060 |
| | 6 | 9am | | | | | | | | | | **0.475** | -0.156 | -0.072 | 0.151 | 0.044 | **0.266** | 0.061 | 0.166 | 0.153 | 0.026 | **0.202** | 0.156 | 0.091 | **0.221** | 0.166 |
| | | 2pm | | | | | | | | | | | **-0.486** | -0.013 | 0.063 | **0.188** | 0.146 | 0.064 | **0.244** | **0.216** | 0.003 | 0.147 | **0.277** | 0.125 | **0.206** | **0.181** |
| | 7 | 9am | | | | | | | | | | | | **0.213** | **0.229** | **0.200** | **0.208** | **0.421** | **0.278** | **0.332** | -0.010 | 0.004 | 0.028 | -0.159 | -0.025 | -0.053 |
| | | 2pm | | | | | | | | | | | | | 0.089 | **0.203** | **0.180** | 0.156 | **0.297** | **0.201** | 0.069 | **0.222** | -0.012 | 0.020 | -0.063 | -0.120 |
| Offset | 5 | 9am | | | | | | | | | | | | | | **0.356** | **0.297** | **0.215** | 0.166 | **0.334** | -0.038 | -0.030 | 0.102 | 0.047 | 0.028 | -0.021 |
| | | 2pm | | | | | | | | | | | | | | | **0.228** | **0.228** | **0.220** | **0.224** | 0.048 | **0.184** | -0.042 | -0.082 | -0.079 | 0.019 |
| | 6 | 9am | | | | | | | | | | | | | | | | **0.334** | **0.426** | **0.392** | -0.077 | 0.043 | **0.187** | 0.015 | 0.015 | 0.035 |
| | | 2pm | | | | | | | | | | | | | | | | | **0.292** | **0.492** | -0.089 | 0.112 | 0.153 | 0.059 | -0.057 | 0.047 |
| | 7 | 9am | | | | | | | | | | | | | | | | | | **0.517** | 0.032 | 0.016 | 0.029 | 0.080 | 0.011 | 0.051 |
| | | 2pm | | | | | | | | | | | | | | | | | | | -0.115 | 0.064 | 0.047 | 0.033 | 0.030 | -0.021 |
| Onset | 5 | 9am | | | | | | | | | | | | | | | | | | | | **0.324** | **0.193** | **0.208** | 0.118 | **0.181** |
| | | 2pm | | | | | | | | | | | | | | | | | | | | | 0.127 | 0.095 | **0.250** | **0.197** |
| | 6 | 9am | | | | | | | | | | | | | | | | | | | | | | **0.278** | **0.377** | **0.231** |
| | | 2pm | | | | | | | | | | | | | | | | | | | | | | | **0.226** | **0.342** |
| | 7 | 9am | | | | | | | | | | | | | | | | | | | | | | | | **0.263** |
| | | 2pm | | | | | | | | | | | | | | | | | | | | | | | | |

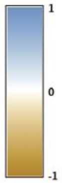

**Supp. Tab. 2:** Table of correlations between the startle responses (tapping, dark flash, onset and offset) and the different conditions (spontaneous, dark intervals and light intervals) for all days and times tested. Values given are the r values from Pearson's correlations, with color scale based on that r value (blue positive correlation and yellow negative correlation). Significant correlations are denoted by the bold font when $p < 0.05$.

| | | | Dark Intervals | | | | | | Light Intervals | | | | | | Spontaneous | | | | | |
|---|---|---|---|---|---|---|---|---|---|---|---|---|---|---|---|---|---|---|---|---|
| | | | 5 | | 6 | | 7 | | 5 | | 6 | | 7 | | 5 | | 6 | | 7 | |
| | | | 9am | 2pm | 9am | 2pm | 9am | 2pm | 9am | 2pm | 9am | 2pm | 9am | 2pm | 9am | 2pm | 9am | 2pm | 9am | 2pm |
| Tapping | 5 | 9am | 0.131 | 0.152 | **0.192** | -0.035 | 0.059 | 0.034 | -0.024 | 0.014 | 0.025 | -0.214 | -0.060 | -0.052 | -0.011 | 0.076 | -0.033 | **-0.186** | 0.038 | -0.138 |
| | | 2pm | -0.063 | 0.032 | 0.026 | **-0.213** | -0.028 | -0.025 | **0.190** | **0.234** | 0.038 | **-0.342** | -0.089 | -0.144 | **0.185** | **0.382** | -0.083 | **-0.208** | -0.076 | -0.166 |
| | 6 | 9am | -0.047 | 0.039 | 0.032 | -0.148 | 0.077 | 0.055 | -0.062 | 0.058 | -0.121 | **-0.322** | -0.063 | -0.002 | 0.112 | 0.153 | 0.037 | **-0.283** | 0.023 | -0.069 |
| | | 2pm | -0.191 | -0.104 | -0.043 | 0.053 | 0.127 | 0.166 | -0.163 | -0.130 | -0.112 | **0.195** | **0.290** | **0.328** | 0.064 | 0.052 | 0.040 | 0.030 | **0.227** | 0.153 |
| | 7 | 9am | -0.110 | 0.012 | 0.035 | **-0.198** | 0.057 | 0.032 | 0.006 | 0.094 | -0.057 | **-0.302** | 0.019 | 0.104 | 0.066 | 0.156 | 0.060 | **-0.297** | 0.038 | -0.109 |
| | | 2pm | -0.151 | -0.078 | **-0.233** | **-0.308** | **-0.176** | **-0.177** | -0.058 | -0.023 | -0.099 | **-0.283** | -0.113 | -0.120 | -0.049 | 0.000 | -0.025 | -0.159 | -0.111 | -0.111 |
| Dark Flash | 5 | 9am | 0.129 | 0.120 | 0.166 | -0.171 | 0.052 | -0.019 | 0.000 | 0.138 | -0.020 | **-0.262** | -0.044 | -0.096 | 0.151 | **0.222** | 0.107 | -0.142 | 0.087 | -0.111 |
| | | 2pm | -0.061 | 0.014 | -0.015 | 0.029 | 0.012 | 0.007 | 0.121 | 0.010 | -0.042 | 0.035 | -0.020 | -0.068 | -0.099 | -0.047 | -0.076 | -0.053 | -0.121 | -0.012 |
| | 6 | 9am | -0.013 | 0.067 | 0.081 | -0.082 | 0.111 | 0.045 | 0.089 | 0.153 | -0.077 | -0.119 | 0.027 | 0.007 | 0.103 | 0.131 | **0.213** | -0.143 | 0.116 | -0.098 |
| | | 2pm | 0.084 | 0.155 | **0.187** | 0.035 | **0.254** | **0.184** | 0.027 | -0.064 | 0.005 | -0.059 | 0.105 | -0.024 | 0.020 | 0.064 | 0.016 | -0.090 | **0.175** | 0.042 |
| | 7 | 9am | 0.045 | -0.016 | 0.130 | **0.204** | **0.202** | 0.099 | -0.023 | -0.149 | -0.073 | **0.263** | 0.165 | 0.030 | -0.143 | -0.061 | 0.063 | **0.284** | **0.260** | 0.177 |
| | | 2pm | 0.160 | 0.080 | -0.072 | -0.026 | 0.133 | 0.061 | -0.167 | -0.112 | -0.030 | -0.030 | 0.051 | -0.004 | -0.108 | -0.135 | -0.052 | 0.107 | 0.127 | 0.093 |
| Offset | 5 | 9am | 0.136 | 0.033 | 0.162 | -0.008 | 0.142 | 0.012 | -0.005 | 0.100 | 0.008 | -0.098 | -0.154 | **-0.326** | 0.166 | 0.202 | **0.179** | 0.090 | -0.022 | **-0.172** |
| | | 2pm | 0.150 | 0.110 | 0.064 | -0.016 | 0.067 | -0.007 | -0.059 | 0.023 | -0.053 | -0.059 | -0.082 | -0.133 | -0.022 | 0.115 | -0.094 | -0.051 | 0.022 | -0.153 |
| | 6 | 9am | -0.005 | 0.012 | 0.157 | -0.001 | **0.203** | 0.109 | 0.021 | -0.060 | -0.078 | -0.008 | 0.009 | -0.186 | -0.080 | 0.090 | **0.217** | -0.096 | 0.131 | -0.042 |
| | | 2pm | -0.048 | -0.063 | 0.017 | 0.143 | **0.224** | 0.123 | -0.011 | **-0.196** | -0.134 | **0.303** | 0.132 | -0.070 | -0.079 | -0.016 | 0.027 | **0.262** | 0.162 | 0.055 |
| | 7 | 9am | 0.002 | -0.067 | 0.053 | 0.016 | **0.285** | **0.200** | 0.061 | -0.042 | -0.078 | 0.084 | **0.200** | 0.023 | 0.058 | -0.020 | 0.026 | -0.011 | **0.270** | 0.140 |
| | | 2pm | -0.098 | -0.085 | 0.042 | 0.089 | **0.345** | **0.244** | 0.007 | -0.087 | -0.066 | **0.193** | **0.186** | -0.081 | 0.143 | 0.126 | 0.150 | 0.159 | **0.251** | **0.193** |
| Onset | 5 | 9am | **0.497** | **0.466** | **0.278** | **0.314** | **0.305** | **0.340** | 0.031 | 0.128 | 0.067 | 0.049 | 0.171 | **0.275** | -0.055 | -0.082 | 0.094 | 0.056 | **0.245** | **0.225** |
| | | 2pm | **0.340** | **0.397** | 0.111 | -0.001 | 0.151 | 0.066 | -0.004 | 0.112 | -0.010 | -0.072 | 0.107 | -0.027 | 0.012 | 0.104 | 0.024 | 0.063 | **0.197** | -0.005 |
| | 6 | 9am | **0.240** | **0.262** | **0.512** | **0.309** | **0.395** | **0.355** | **0.172** | 0.067 | -0.022 | 0.111 | 0.056 | -0.015 | 0.032 | 0.070 | **0.218** | 0.027 | 0.114 | 0.073 |
| | | 2pm | 0.110 | 0.110 | 0.103 | **0.317** | **0.204** | **0.244** | 0.008 | -0.009 | -0.085 | 0.114 | -0.095 | 0.079 | -0.071 | -0.135 | 0.056 | 0.064 | 0.031 | 0.152 |
| | 7 | 9am | 0.119 | **0.228** | **0.314** | 0.105 | **0.364** | **0.282** | 0.079 | 0.136 | 0.007 | -0.020 | 0.103 | -0.013 | **0.189** | **0.193** | **0.214** | -0.046 | **0.188** | -0.056 |
| | | 2pm | -0.058 | -0.051 | 0.127 | 0.089 | **0.186** | 0.153 | -0.025 | -0.048 | -0.024 | 0.040 | 0.039 | 0.093 | -0.027 | -0.022 | 0.136 | 0.011 | 0.066 | 0.007 |

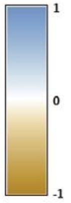

**Supp. Tab. 4:** Table of correlations between days for each morphological parameter (resting heart rate (BPM; beats per minute), body length and swim bladder size). All correlation coefficients were derived from calculating Pearson's correlations, with color scale based on that r value (blue positive correlation and yellow negative correlation). Significant correlations are denoted by the bold font when $p < 0.05$.

| Morphological Parameter | DPF Comparison | r | p value |
|---|---|---|---|
| BPM | 5 v 6 | **0.48** | **4.41** $\times 10^{-9}$ |
| | 5 v 7 | **0.61** | **1.50** $\times 10^{-14}$ |
| Body Length | 5 v 6 | **0.79** | **4.96** $\times 10^{-29}$ |
| | 5 v 7 | **0.64** | **2.16** $\times 10^{-16}$ |
| Swim Bladder Size | 5 v 6 | **0.63** | **1.06** $\times 10^{-15}$ |
| | 5 v 7 | **0.32** | **1.97** $\times 10^{-4}$ |

**Supp. Tab. 5:** Table of correlations between each morphological parameter (resting heart rate (BPM; beats per minute), body length and swim bladder size) and the startle responses (tapping, dark flash, offset and onset). All correlation coefficients were derived from calculating Pearson's correlations, with color scale based on that r value (blue positive correlation and yellow negative correlation). Significant correlations are denoted by the bold font when $p < 0.05$.

| Morphological Parameter | DPF | Tapping | | Dark Flash | | Offset | | Onset | |
|---|---|---|---|---|---|---|---|---|---|
| | | r | p value | r | p value | r | p value | r | p value |
| BPM | 5 | **0.409** | **1.11** $\times10^{-6}$ | 0.109 | 0.214 | -0.004 | 0.963 | 0.030 | 0.734 |
| | 6 | 0.152 | 0.083 | **0.242** | **0.005** | -0.071 | 0.418 | 0.103 | 0.238 |
| | 7 | **0.435** | **1.93** $\times10^{-7}$ | -0.122 | 0.163 | -0.006 | 0.942 | **0.209** | **0.016** |
| Body Length | 5 | **0.323** | **1.57** $\times10^{-4}$ | 0.155 | 0.077 | **0.214** | **0.014** | 0.046 | 0.604 |
| | 6 | 0.074 | 0.397 | **0.211** | **0.015** | **0.240** | **0.005** | 0.074 | 0.401 |
| | 7 | 0.015 | 0.865 | **0.322** | **1.71** $\times10^{-4}$ | **0.379** | **7.4** $\times10^{-6}$ | 0.123 | 0.160 |
| Swim Bladder Size | 5 | -0.148 | 0.090 | **-0.192** | **0.027** | -0.124 | 0.157 | 0.099 | 0.259 |
| | 6 | **-0.361** | **2.16** $\times10^{-5}$ | **-0.430** | **2.60** $\times10^{-6}$ | -0.014 | 0.874 | -0.061 | 0.488 |
| | 7 | **-0.501** | **9.70** $\times10^{-10}$ | 0.161 | 0.066 | 0.026 | 0.769 | -0.106 | 0.288 |
